# *Cis* and *Trans* Regulatory Mechanisms of ecDNA Segregation

**DOI:** 10.1101/2024.12.31.630921

**Authors:** Yipeng Xie, Jun Yi Stanley Lim, Wenyue Liu, Collin Gilbreath, Xiaohui Sun, Kailiang Qiao, Yoon Jung Kim, Sihan Wu

## Abstract

Extrachromosomal DNAs (ecDNAs) attach to chromosomes during mitosis for random segregation and promote cancer heterogeneity. However, the mechanism governing ecDNA-chromosome mitotic interactions remains poorly understood. This study shows that ecDNAs tether to histone H3 lysine 27 acetylation (H3K27ac)-marked chromatin during mitosis. Depleting H3K27ac disrupts this interaction. Diverse bromodomain proteins, as H3K27ac readers, stabilise ecDNA-chromosome binding in a context-dependent and complementary manner. Although disrupting the Mediator complex in asynchronous cells detaches ecDNAs from mitotic chromosomes, Mediator and active Pol II are absent from ecDNAs during mitosis, suggesting that ecDNAs are transcriptionally silent during mitosis. Instead, inactive Pol II mediates ecDNA attachment. Furthermore, CRISPR interference targeting transcriptional regulatory elements on ecDNA impairs ecDNA segregation. Mis-segregated ecDNAs were expelled into the cytosol, leading to diminished oncogene expression and a reversal of therapy resistance. Our research provides universal *cis* and *trans* regulatory mechanisms of ecDNA segregation, offering deeper insight into ecDNA-driven oncogenesis.

## Introduction

DNA transmission to daughter cells during cell division is a fundamental process in inheritance, ensuring the stability of genetic information across generations.^1^ In eukaryotes, chromosomal inheritance involves spindle fibre attachment to the kinetochore at the centromere, which facilitates the equal distribution of chromosomes between two daughter cells.^2,3^ However, the inheritance mechanism for acentric extrachromosomal DNA (ecDNA), a prevalent oncogene carrier in aggressive tumours, remains unclear.

ecDNA is chromatinized circular DNA specifically found in cancer with a typical size ranging from 50-kb to 5-Mb.^4^ It is present in nearly half of all human cancer types and up to one-third of cancer patients throughout their disease.^5–7^ ecDNA carries diverse oncogenes, immunoregulatory genes, and associated enhancers.^8,9^ Owing to high copy numbers and hyper-accessible chromatin, ecDNA can drive massive oncogene expression.^10^ More importantly, a lack of functional centromeres causes ecDNAs to segregate randomly during cancer cell division, similar to plasmid inheritance in bacteria.^11–15^ This random distribution generates a heterogeneous pool of cancer cells with varying oncogene copy numbers in every cell cycle. In several cancer types, including small cell lung cancer, glioblastoma, and medulloblastoma, ecDNA’s asymmetric segregation has been shown to promote rapid tumour evolution, stress adaptation, and therapy resistance.^16–18^ Additionally, a recent study has shown that distinct ecDNA species within a single cell can co-segregate into daughter cells.^19^ Such behaviour further enables ecDNA to evade the principles of Mendelian inheritance and foster tumour aggressiveness. Thus, revealing the molecular mechanism mediating ecDNA segregation is critical for understanding cancer genome maintenance and developing effective therapeutic strategies for ecDNA-driven cancers.

How do ecDNAs segregate during mitosis? Historically, it was proposed that ecDNAs attach to chromosomes for segregation as early as 1978.^20^ Subsequent imaging studies have confirmed this finding, showing that ecDNAs hitchhike on mitotic chromosomes during cell division.^13,21,22^ A recent study suggests that BRD4 and mitotic transcription activity regulate *MYC* ecDNAs segregation.^23^ However, the generalised mechanism underlying the segregation of a broad spectrum of ecDNA species remains to be elucidated.

The attachment of ecDNAs to mitotic chromosomes represents a unique form of chromatin-chromatin interaction. Electrostatic repulsion between DNA molecules, arising from their negatively charged phosphate backbones, necessitates the involvement of local chromatin modifications and DNA-binding proteins to enable ecDNA-chromosome interactions during mitosis. The present study aims to investigate the histone modifications and protein complexes that facilitate ecDNA segregation, thereby providing new insights into the mechanism of oncogenic inheritance in cancer.

## Results

### 1. ecDNA *trans*-interaction sites on mitotic chromosomes are enriched with H3K27ac and H3K4me3 chromatin

We previously demonstrated that, even during metaphase, ecDNA chromatin is enriched with active histone modifications (Extended Data Fig. 1a), including H3K27 acetylation (H3K27ac) and H3K4 tri-methylation (H3K4me3).^10^ In addition, ecDNAs may interact with chromosomes via their active chromatin in an enhancer-promoter contact fashion.^24^ This prompted us to investigate whether H3K27ac and H3K4me3, which mark active enhancers and promoters, are required for ecDNA to interact with mitotic chromosomes. We used the MSTO211H lung cancer cell line as our model system and performed *in situ* HiC to profile ecDNA *trans*-interaction sites during mitosis and correlated them with H3K27ac and H3K4me3 ChIP-seq data (Fig. 1a, Extended Data Fig. 1b). Mitotic cells were enriched with nocodazole treatment and purified by flow cytometric sorting for the phosphorylated MPM2-positive and 4N DNA content fraction. The MPM2-negative, 2N DNA content fraction, representing the G1 phase cells, was sorted as control. The MSTO211H cell line carries *MYC* ecDNAs made of a simple structure originating from a 500-kb DNA segment of chromosome 8, enabling us to distinguish ecDNA *trans*-interaction from chromosome translocation and ecDNA rearrangement (Extended Data Fig. 1c-d).

**Fig. 1.**
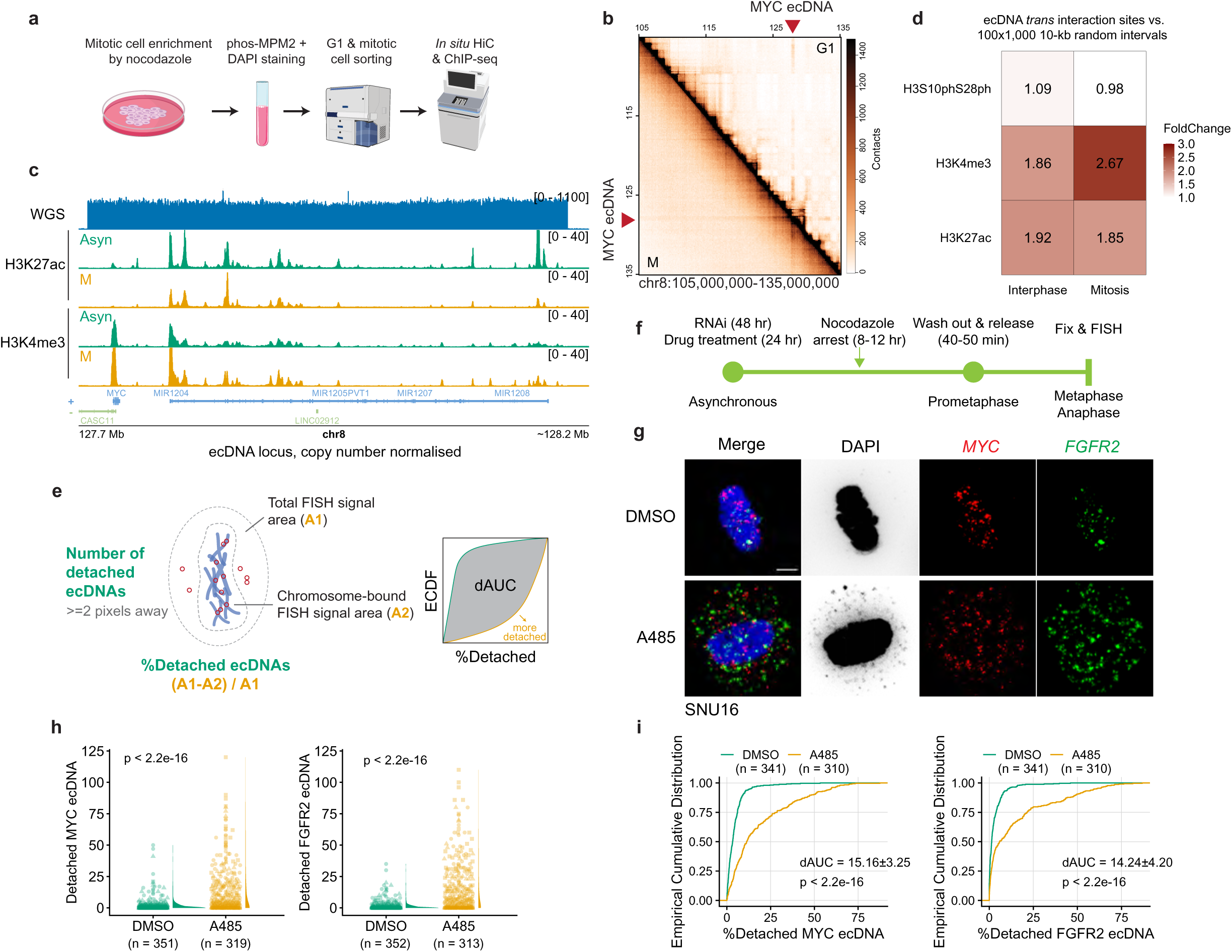
Epigenetic regulators of ecDNA segregation. **(a)** Illustration of profiling ecDNA *trans*-interaction sites between G1 and M phases. **(b)** HiC contact heatmap showing *MYC* ecDNA interactions between G1 and M phases in MSTO211H cells. The *MYC* ecDNA locus is highlighted. **(c)** ChIP-seq coverage tracks for H3K27ac and H3K4me3 in asynchronous (Asyn) or mitotic (M) MSTO211H cells. **(d)** Heatmap of H3S10phS28ph, H3K27ac, and H3K4me3 coverage comparison between ecDNA *trans*-interaction sites and 100 iterations of 1,000 randomly selected 10-kb genomic intervals in interphase and mitotic MSTO211H cells. Each box represents the sequencing coverage of ecDNA *trans* interaction sites normalized to random genomic intervals. **(e)** Quantification strategy of detached ecDNAs during mitosis. Both the number of detached ecDNAs and the percentage of detached ecDNAs out of total ecDNAs were quantified. **(f)** Workflow showing the imaging approach to study ecDNA interaction with chromosomes during mitosis. An 8 to 12-hr nocodazole treatment (unless specified) was applied to enrich prometaphase cells after drug treatment or RNAi transfection, followed by a washout to release the cell cycle into the metaphase and anaphase. **(g)** Representative *MYC* and *FGFR2* ecDNA FISH on 10 μM A485 treated SNU16 metaphase cells (scale bar: 5 μm). **(h-i)** The number **(h)** and percentage **(i)** of detached *MYC* and *FGFR2* ecDNA per SNU16 metaphase cell after A485 treatment (two-sided Wilcoxon test). Nocodazole treatment: 12-16 hr. Data were pooled from three replicates for statistics. dAUC was shown as mean ± SEM.

Over 600 million valid HiC interaction pairs per library were obtained, with high reproducibility between replicates (Extended Data Fig. 1e-f). Compared to the G1 phase, topologically associating domains in M phase cells were largely reduced in agreement with previous reports^25^ (Fig. 1b, Extended Data Fig. 1g). ecDNA-derived global *trans*-interactions were also detected as previously observed,^24^ but notably decreased at the M phase (Fig. 1b, Extended Data Fig. 1g), suggesting fewer regions from mitotic chromosomes remained accessible for ecDNA to interact with.

To analyse ecDNA *trans*-interaction sites, HiC replicates were merged, and *trans*-interactions were called at a 10-kb resolution. Significant *trans*-interactions were defined as the top 0.1% of ecDNA contacts with non-ecDNA-originated chromosomes, excluding chromosome 8, where *MYC* ecDNAs originated (Extended Data Fig. 1h), yielding 584 significant ecDNA *trans*-interaction pairs during mitosis. The number of unique mitotic *trans*-interaction sites was not correlated with the chromosome size, suggesting a non-random ecDNA attachment mechanism (Extended Data Fig. 1i).

In parallel, ChIP-seq analysis showed that genome-wide H3K27ac modification was profoundly decreased during mitosis, whereas H3K4me3 was retained, consistent with previous reports^26^ (Extended Data Fig. 1j-k). In contrast, H3K27ac on ecDNA remained. Compared with chromosomes, ecDNAs showed higher H3K27ac levels during mitosis, consistent with the immunofluorescence staining results (Fig. 1c, Extended Data Fig. 1a & 1l).

Next, we analysed the association of ecDNA *trans*-interaction sites with active histone marks. Compared with 100 iterations of 1,000 randomly selected 10-kb genomic intervals without duplicated sampling, ecDNA *trans*-interaction sites during interphase were enriched with H3K27ac and H3K4me3, concordant with the previous study^24^ (Fig. 1d). This pattern remained in mitotic cells, suggesting a shared mechanism of ecDNA *trans*-interaction between interphase and mitosis (Fig. 1d, Extended Data Fig. 1m). In contrast, the phosphorylation of H3 at serine 10 and 28, a histone modification that is globally deposited during mitosis but less abundant during interphase,^27^ did not show an enrichment, further supporting the specificity for H3K27ac and H3K4me3 (Fig. 1d, Extended Data Fig 1n). Together, these data indicate that ecDNAs may be preferentially associated with active chromatin regions during segregation.

### 2. H3K27ac is important for ecDNA attachment to mitotic chromosomes

H3K27ac is mainly enriched at active enhancers and promoters. We applied *in situ* DNA FISH imaging and quantification to study the attachment of ecDNAs to mitotic chromosomes. Two orthogonal metrics were employed to assess ecDNA detachment per cell (Fig. 1e): (1) the number of detached ecDNAs, and (2) the proportion of detached ecDNAs. The latter enables the estimation of the average effect size by calculating the difference in the area under the curve (dAUC), derived from the empirical cumulative distribution function, where dAUC ranges from 0 to 100, representing 0% to 100% detachment. In COLO320HSR cells, where all ecDNAs were reinserted into chromosomes, *MYC* DNA FISH imaging served as a negative control to establish the threshold for identifying detached ecDNAs, defined as signals located at least two pixels away from mitotic chromosomes. This criterion effectively eliminated false positive signals and was sufficient to identify detached ecDNAs based on our imaging resolution (Extended Data Fig. 2a-c).

A485, the CBP/P300 inhibitor,^28^ depleted global H3K27ac as measured by CUT&RUN and Western blot in MSTO211H cells (Fig. 1f, Extended Data Fig. 2d-g). Concomitantly, about 16.87% of *MYC* ecDNAs were significantly detached from chromosomes during metaphase as determined by dAUC (Extended Data Fig. 2h-j). More than 44.57% of cells exhibited at least 20% ecDNA detachment within one cell cycle upon A485 treatment, representing a 5.2-fold increase compared with the control (Extended Data Fig. 2j). The accuracy of dAUC estimation was orthogonally validated by qPCR-based quantification of ecDNA copy number. Treatment with 2 μM A485 reduced *MYC* ecDNA copy number from 90.8 to 10.3 over 3 weeks with a cell doubling time of 35.6 hours, corresponding to a 14.9% loss per cell cycle, closely aligning with imaging-based estimates (Extended Data Fig. 2k).

We noticed cell-to-cell variability in ecDNA detachment rates. To address this heterogeneity, we performed H3K27ac immunofluorescence in combination with *MYC* DNA FISH in A485-treated cells. Although H3K27ac levels were globally reduced upon A485 treatment, cells with <5% ecDNA detachment retained significantly higher residual H3K27ac signal on ecDNAs than those with >30% ecDNA detachment (Extended Data Fig. 2l-m). This result further reinforced supporting the importance of H3K27ac in ecDNA mitotic tethering.

The effect of A485 treatment was independently verified in SNU16 stomach cancer cells with *MYC* and *FGFR2* ecDNAs (Fig. 1g-i, Extended Data Fig. 2n) and COLO320DM colon cancer cells with *MYC* ecDNAs (Extended Data Fig. 2o-r), suggesting that H3K27ac is commonly required for ecDNA interaction with mitotic chromosomes. This phenomenon was not confounded by nocodazole-induced mitotic cell enrichment, as A485 treatment also detached *MYC* and *FGFR2* ecDNAs in natural mitotic cells, while nocodazole treatment alone had no effects (Extended Data Fig. 2s-t).

Two strategies further validated the involvement of H3K27ac. First, we used a histone deacetylase inhibitor, SAHA, to counteract the effect of A485. In MSTO211H cells, SAHA treatment restored H3K27ac modification suppressed by A485 (Extended Data Fig. 2d-g). In the meantime, *MYC* ecDNAs’ attachment to mitotic chromosomes was also recovered (Extended Data Fig. 2h-j). RNAi knockdown of *CBP* (*CREBBP*) and *P300* (*EP300*), the writers of H3K27ac, resulted in the same ecDNA detachment phenotype as the A485 treatment (Extended Data Fig. 3a-c). These data collectively show that H3K27ac is an important histone modification for ecDNA attachment to mitotic chromosomes.

H3K4 methylation is deposited by the COMPASS complex, which contains diverse methyltransferases from the SET1 and MLL families and shared subunits, including DPY30 and RBBP5^29^ (Extended Data Fig. 2d). Global H3K4me3 level was downregulated when *DPY30* and *RBBP5* were knocked down by RNAi (Extended Data Fig. 3d). However, neither *MYC* nor *FGFR2* ecDNAs were significantly detached from mitotic chromosomes in SNU16 cells (Extended Data Fig. 3e-f), suggesting that H3K4me3 is not a key player in ecDNA segregation.

### 3. Diverse bromodomain proteins anchor ecDNAs to mitotic chromosomes

While H3K27ac is shown to facilitate ecDNA attachment, it does not fully explain why ecDNA-chromosome interaction becomes feasible, as the acetyl group is insufficient to neutralise the negative charge of the DNA backbone. Thus, we hypothesised that proteins bound to H3K27ac chromatin are necessary to mediate ecDNAs’ mitotic interaction with chromosomes.

Bromodomain-containing proteins are readers of H3K27ac chromatin. A previous study demonstrates that BRD4 facilitates *MYC* ecDNA segregation.^23^ We tested this argument by using dBET6, a PROTAC molecule targeting the bromodomain and extraterminal (BET) proteins,^30^ including BRD4, in MSTO211H cells (Extended Data Fig. 4a-b). However, contrasting results were observed depending on the treatment regimen (Fig. 2a). Only when cells first entered mitosis, but not before, could dBET6 significantly detach *MYC* ecDNAs (Fig. 2b-d).

**Fig. 2.**
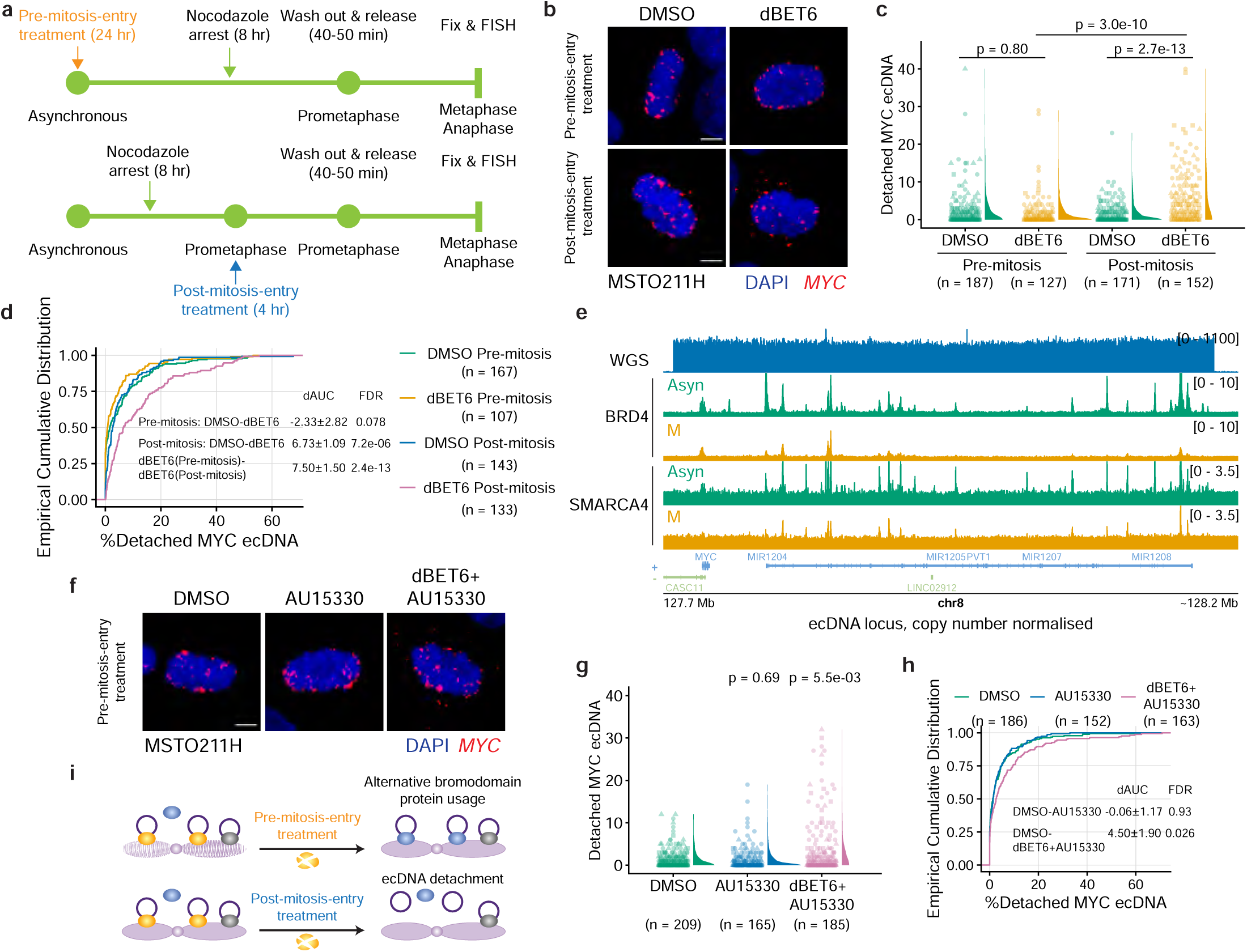
Bromodomain proteins anchor ecDNAs to mitotic chromosomes. **(a)** Treatment strategies to manipulate bromodomain proteins. Cells were treated either with pre-mitosis-entry arrest or post-mitosis-entry arrest. **(b)** Representative *MYC* ecDNA FISH on 200 nM dBET6 (pre-mitosis-entry) or 1 μM dBET6 (post-mitosis-entry) treated MSTO211H metaphase cells (scale bar: 5 μm). **(c-d)** The number **(c)** and percentage **(d)** of detached *MYC* ecDNAs per MSTO211H metaphase cell after pre-mitosis-entry or post-mitosis-entry dBET6 treatment (Kruskal-Wallis test with post-hoc Dunn test). Nocodazole treatment: 8 hr. Data were pooled from three replicates for statistics. dAUC was shown as mean ± SEM. **(e)** ChIP-seq coverage tracks for BRD4 and SMARCA4 in asynchronous (Asyn) or mitotic (M) MSTO211H cells. **(f)** Representative *MYC* ecDNA FISH on 1 μM AU15330 or 200 nM dBET6+1 μM AU15330 treated (pre-mitosis-entry) MSTO211H metaphase cells (scale bar: 5 μm). **(g-h)** The number **(g)** and percentage **(h)** of detached *MYC* ecDNAs per MSTO211H metaphase cell after AU15330 or dBET6+AU15330 pre-mitosis-entry treatment (Kruskal-Wallis test with post-hoc Dunn test). Nocodazole treatment: 8 hr. Data were pooled from three replicates for statistics. dAUC was shown as mean ± SEM. **(i)** Proposed mechanisms of how BET proteins and SWI/SNF proteins mediate ecDNA attachment.

The phenotypic contrast associated with the treatment regimen led us to hypothesise that chronic depletion of selected bromodomain proteins in interphase would induce alternative usage of bromodomain proteins to establish ecDNA-chromosome interactions. Transcriptomic analysis of human bromodomain proteins from Uniprot in common ecDNA-containing cancer cell lines^10,31^ showed that the most highly expressed genes belong to two families: the BET proteins, including BRD2, BRD3, and BRD4, and the SWI/SNF complexes, including SMARCA2/4 and BRD7/9 (Extended Data Fig. 4c). This prompted us to study whether SWI/SNF proteins function as alternative anchors for ecDNA attachment.

First, ChIP-seq analysis in MSTO211H cells showed that both BRD4 and SMARCA4 remained associated with *MYC* ecDNAs during mitosis, despite a notable reduction compared to asynchronous cells (Fig. 2e). Second, although dBET6 treatment effectively depleted BRD4 binding to *MYC* ecDNAs, SMARCA4 association persisted (Extended Data Fig. 4d). Conversely, treatment with AU15330, a SMARCA2/4 PROTAC molecule,^32^ diminished SMARCA4 binding to ecDNAs, while BRD4 remained bound. These data suggest functional redundancy among these bromodomain proteins (Extended Data Fig. 4d-e).

Consistent with these findings, AU15330 treatment alone before mitotic entry did not significantly disrupt *MYC* ecDNA attachment in MSTO211H cells, similar to dBET6 treatment alone (Fig. 2b-d). However, it synergised with dBET6 to exacerbate ecDNA detachment from mitotic chromosomes (Fig. 2f-h). These results indicate that multiple bromodomain proteins participate in the interaction between ecDNA and mitotic chromosomes.

This mechanism applies to other ecDNA species. In SNU16 cells harbouring *MYC* and *FGFR2* ecDNAs, post-mitosis-entry treatment of dBET6 led to more ecDNA detachment compared to pre-mitosis-entry treatment (Extended Data Fig. 5a-c). Similarly, co-treatment with dBET6 and AU15330 before mitotic entry detached more ecDNAs compared to AU15330 alone (Extended Data Fig. 5d-f).

Collectively, these results suggest that diverse bromodomain proteins can act interchangeably during interphase to establish ecDNA-chromosome interactions, which can compensate for the loss of individual proteins, priming ecDNA for mitotic attachment. However, upon mitotic entry, the use of bromodomain proteins is committed within each ecDNA molecule, limiting the flexibility for alternative bromodomain protein usage (Fig. 2i).

### 4. The Mediator complex primes ecDNA attachment in interphase

BET and SWI/SNF proteins are critical regulators of enhancer-promoter interactions, facilitating the assembly of the transcription machinery through their interactions with the Mediator complex.^33,34^ We then investigated whether the Mediator is also involved in ecDNA segregation.

The core Mediator is a large, multi-subunit complex whose functional activity is regulated by its kinase module, which includes cyclin-dependent kinases, notably CDK8 (Fig. 3a).^35^ The knockdown of *CDK8* significantly detached ecDNAs from mitotic chromosomes in MSTO211H and SNU16 cells (Fig. 3b-d, Extended Data Fig. 6a-d), suggesting a pivotal role for the Mediator complex in the segregation of ecDNAs. The dependency on CDK8’s kinase activity was confirmed by Cyclin C knockdown, an essential regulator for CDK8 activation, which similarly resulted in the detachment of ecDNAs in both cell lines (Extended Data Fig. 6a-g).

**Fig. 3.**
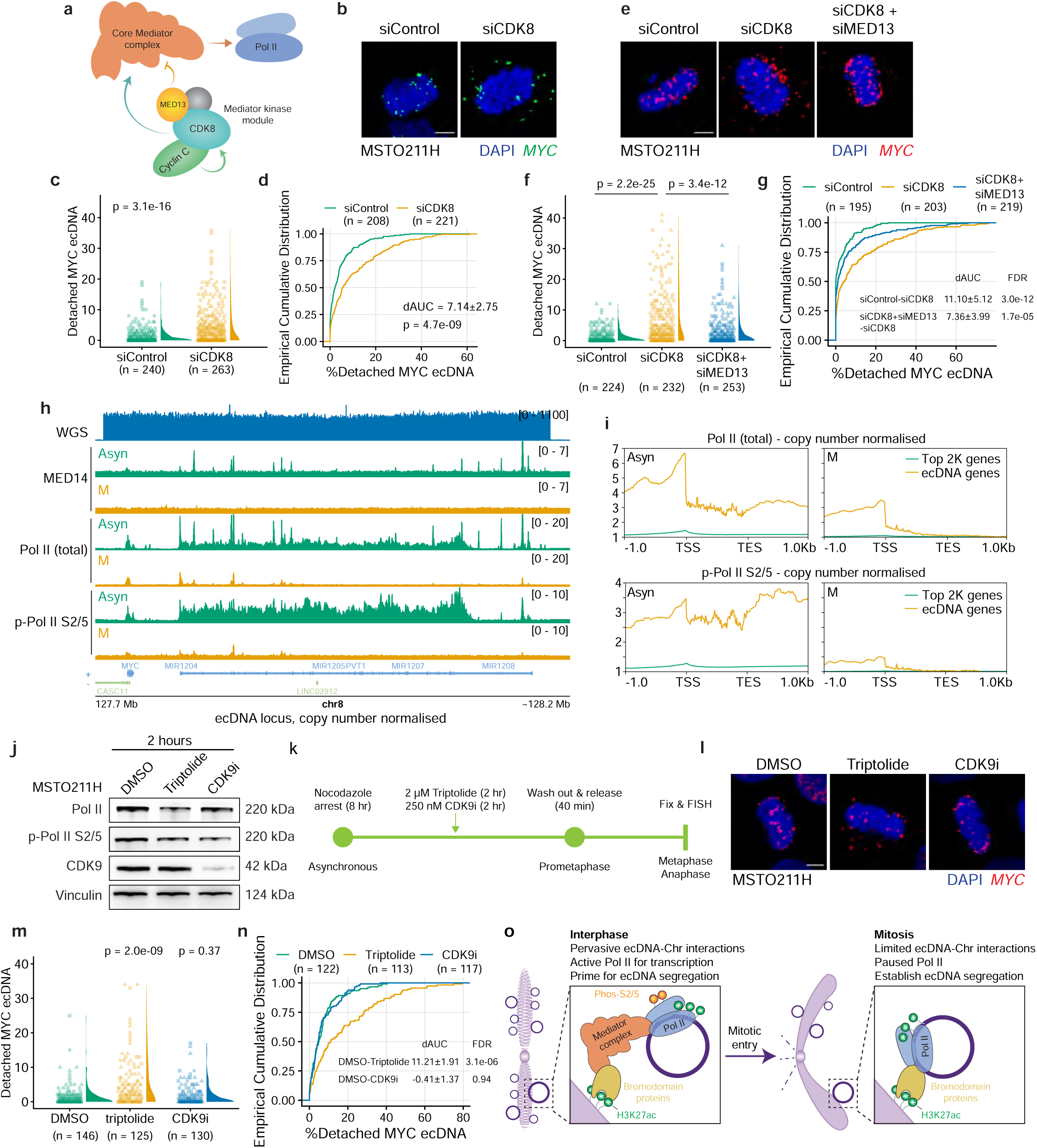
The Mediator complex and RNA polymerase II regulate ecDNA segregation. **(a)** Illustration depicting the Mediator complex components. **(b)** Representative *MYC* ecDNA FISH on MSTO211H metaphase cells after *CDK8* knockdown (scale bar: 5 μm). **(c-d)** The number **(c)** and percentage **(d)** of detached *MYC* ecDNAs per MSTO211H metaphase cell after *CDK8* RNAi knockdown (two-sided Wilcoxon test). Nocodazole treatment: 8-12 hr. Data were pooled from three replicates for statistics. dAUC was shown as mean ± SEM. **(e)** Representative *MYC* ecDNA FISH images showing the rescue effect of *MED13* knockdown in ecDNA attachment to mitotic chromosomes (scale bar: 5 μm). **(f-g)** The number **(f)** and percentage **(g)** of detached *MYC* ecDNAs per MSTO211H metaphase cell after *CDK8* and *MED13* RNAi knockdown (Kruskal-Wallis test with post-hoc Dunn test). Nocodazole treatment: 8-12 hr. Data were pooled from three replicates for statistics. dAUC was shown as mean ± SEM. **(h)** ChIP-seq coverage tracks for MED14, Pol II, and p-Pol II Ser2/5 on ecDNA in asynchronous (Asyn) or mitotic (M) MSTO211H cells. **(i)** Quantification for Pol II, and p-Pol II Ser2/5 ChIP-seq read counts on ecDNA versus the top 2,000 expressed genes in asynchronous (Asyn) or mitotic (M) MSTO211H cells. **(j)** Western blot for Pol II, p-Pol II Ser2/5, and CDK9 in MSTO211H cells after 2 μM triptolide or 250 nM CDK9i treatment. **(k)** Workflow showing the imaging approach to study ecDNA attachment with mitotic chromosomes during metaphase. An 8-hr nocodazole treatment was first applied to enrich prometaphase cells. Two hr before the nocodazole treatment endpoint, a transcription inhibitor was added to the cell culture, before washing out to release cells into metaphase and anaphase for fixation and imaging. **(l)** Representative *MYC* ecDNA FISH on 2 μM triptolide or 250 nM CDK9i treated MSTO211H metaphase cells (scale bar: 5 μm). **(m-n)** The number **(m)** and percentage **(n)** of detached *MYC* ecDNAs per MSTO211H metaphase cell after triptolide or CDK9i treatment (Kruskal-Wallis test with post-hoc Dunn test). Nocodazole treatment: 8 hr. Data were pooled from three replicates for statistics. dAUC was shown as mean ± SEM. **(o)** Proposed molecular machinery anchoring ecDNAs to mitotic chromosomes for segregation.

We further investigated the functional engagement of Mediator. The core Mediator complex lacks transcriptional competence until CDK8 activation.^33,36^ Degradation of MED13, another key subunit of the kinase module, liberates the CDK8-containing kinase module from the core Mediator complex. This facilitates the interaction between the Mediator and the RNA Polymerase II preinitiation complex, which is essential for transcription initiation.^33^ Based on this mechanism, we hypothesised that the absence of MED13 would enable ecDNAs to interact with chromosomes independently of CDK8. Upon knocking down *MED13*, ecDNA detachment caused by *CDK8* knockdown was significantly rescued (Fig. 3e-g, Extended Data Fig. 6a). These results indicate that the Mediator complex contributes to ecDNA attachment.

However, ChIP-seq analysis revealed that, while the core Mediator complex, as exemplified by MED14,^37^ was associated with *MYC* ecDNAs in MSTO211H cells in asynchronous cells, it was completely absent from mitotic ecDNAs (Fig. 3h). This finding suggests that the Mediator complex facilitates the establishment of ecDNA-chromosome interactions during interphase but does not serve as a physical tether during mitosis.

### 5. Inactive RNA Pol II couples ecDNAs to mitotic chromosomes

The involvement of H3K27ac modification, bromodomain proteins, and Mediator suggests that components of the transcription machinery may be involved in ecDNA segregation. However, the absence of Mediator binding to mitotic ecDNAs indicates that the transcription preinitiation complex cannot be assembled on ecDNAs during mitosis.^37^ This observation raises two possibilities: (1) ecDNAs are transcriptionally inactive during mitosis, or (2) transcription elongation may persist on ecDNAs independently of Mediator.

To test these hypotheses, we performed ChIP-seq analysis. In MSTO211H cells, RNA polymerase II (Pol II) remained associated with *MYC* ecDNAs during mitosis, although its binding was markedly reduced compared to asynchronous cells (Fig. 3h). Notably, ecDNA loci retained higher Pol II occupancy than the top 2,000 expressed gene loci even during mitosis (Fig. 3i). However, the distribution of Pol II was altered on ecDNAs, with binding predominantly restricted to transcription start sites (TSS) and notably reduced across gene bodies (Fig. 3i). Concordantly, the active form of Pol II, as exemplified by its serine 2/5-phosphorylation (p-Pol II S2/5), essential for transcription initiation and elongation,^38^ was nearly diminished from mitotic ecDNAs, showing only trace levels at TSS on ecDNAs (Fig. 3h-i). Furthermore, 5-ethynyl uridine (5-EU) labelling confirmed the absence of nascent RNA transcripts from ecDNAs during mitosis (Extended Data Fig. 7a). These data collectively show that ecDNAs are likely transcriptionally silent during mitosis.

A previous study has shown that inhibiting transcription activity by triptolide can disrupt *MYC* ecDNA segregation.^23^ We confirmed this finding using a panel of transcription inhibitors, including triptolide, actinomycin D, and α-amanitin, which all effectively suppressed *MYC* transcription and detached *MYC* ecDNAs from mitotic chromosomes after mitotic entry (Extended Data Fig. 7b-f). However, these compounds not only suppressed p-Pol II S2/5 but also degraded total Pol II (Extended Data Fig. 7b),^39^ which could not distinguish whether ecDNA detachment was due to the loss of transcriptional activity or the absence of Pol II itself.

To address this, we employed a PROTAC molecule targeting CDK9 (CDK9i),^40^ which selectively inhibits transcription elongation by blocking the phosphorylation of Pol II S2/5 without affecting the stability of total Pol II (Extended Data Fig. 7g). Unlike triptolide, post-mitosis-entry treatment of CDK9i did not detach ecDNAs from mitotic chromosomes (Fig. 3j-n), suggesting that the presence of inactive Pol II, rather than the transcription activity, is critical for ecDNA attachment during mitosis. Supporting this finding, depletion of cellular RNA with RNase A in mitotic cells did not detach ecDNAs, indicating that RNA transcripts are not required for ecDNA segregation (Extended Data Fig. 7h-m).

Taken together, our study proposes a revised, universal regulatory mechanism underlying ecDNA segregation (Fig. 3o). During interphase, ecDNAs preferentially interact with active chromatin on chromosomes marked by H3K27ac, consistent with previous findings.^24^ These regions recruit diverse bromodomain proteins, which serve as scaffolds for assembling transcription co-activators and machinery, including the Mediator complex and Pol II, in alignment with current domain knowledge.^41^ These interactions can prime ecDNAs for subsequent attachment to chromosomes during mitosis. Upon mitotic entry, chromosomes are condensed, with limited H3K27ac-marked regions remaining accessible for ecDNA attachment. Concurrently, transcriptional activity on ecDNAs is silenced. Inactive Pol II, stalled at the TSS on ecDNAs, together with bromodomain proteins, functions as a physical bridge anchoring ecDNAs to H3K27ac-marked mitotic chromosome regions.

### 6. Transcriptional regulatory elements facilitate ecDNA anchoring to mitotic chromosomes

In mitotic cells, the binding patterns among H3K27ac, BRD4, SMARCA4, and Pol II were similar on ecDNAs, with enrichment primarily at the TSS and upstream regions (Fig. 4a, Extended Data Fig. 8a). This prompted us to investigate whether these transcriptional regulatory DNA elements might contribute to ecDNA attachment during mitosis.

**Fig. 4.**
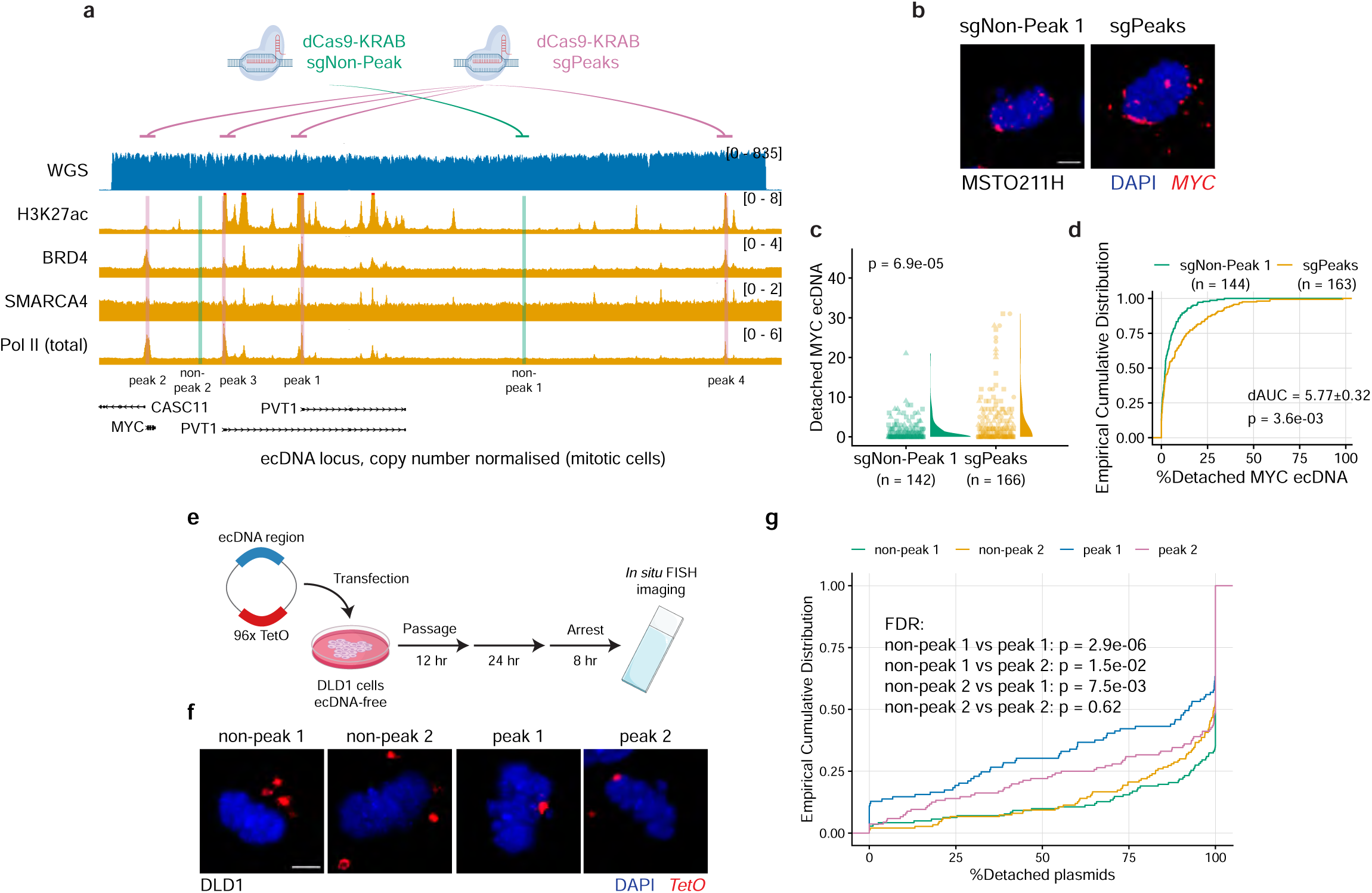
Transcriptionally active ecDNA regions mediate ecDNA attachment. **(a)** ChIP-seq coverage tracks for H3K27ac, BRD4, SMARCA4, and Pol II during mitosis in MSTO211H cells. The chosen “non-peak” and “peak” ecDNA regions for CRISPRi and engineered plasmid construction were labeled in green and pink, respectively. **(b)** Representative *MYC* ecDNA FISH on CRISPRi-treated MSTO211H metaphase cells (scale bar: 5 μm). **(c-d)** The number **(c)** and percentage **(d)** of detached *MYC* ecDNA per MSTO211H metaphase cell after CRISPRi (two-sided Wilcoxon test). Nocodazole treatment: 8 hr. Data were pooled from three replicates for statistics. dAUC was shown as mean ± SEM. **(e)** Workflow showing the imaging approach to study plasmids’ interaction with chromosomes during metaphase. An 8-hr nocodazole treatment was applied to enrich prometaphase cells after transient transfection and one passage, followed by a washout to release the cell cycle into metaphase and anaphase. **(f)** Representative *TetO* plasmid FISH on DLD1 metaphase cells after transient transfection (scale bar: 5 μm). **(g)** Increased percentage of plasmids attached to mitotic chromosomes when the plasmids carried peak regions from ecDNA (Kruskal-Wallis test with post-hoc Dunn test). Nocodazole treatment: 8 hr. Data were pooled from three replicates for statistics.

To test this hypothesis, we employed dCas9-KRAB-mediated CRISPR interference (CRISPRi) to target selected ChIP-seq peak regions on *MYC* ecDNAs in MSTO211H cells, including the *MYC* and *PVT1* promoters. The ecDNA detachment effect increased gradually as the number of targeted peaks rose from one to four (Extended Data Fig. 8b-d). Although the high-level amplification of ecDNAs diluted the CRISPRi efficiency (Extended Data Fig. 8c), and not all peaks on ecDNAs were targeted, targeting four peaks simultaneously successfully detached 5.83% of ecDNAs within a cell cycle compared to CRISPRi targeting a non-peak region (Fig. 4b-d).

To validate this finding using an orthogonal approach, we engineered imaging plasmids containing 96× TetO repeats. The mammalian expression cassette was replaced with a 3-kb DNA segment derived from either peak or non-peak regions (Fig. 4e, Extended Data Fig. 8e). These constructs were transfected into an ecDNA-free, near-diploid cell line, DLD1. After one passage to eliminate residual, extracellular plasmids, cells were cultured for one additional cell cycle to ensure plasmid incorporation (Fig. 4e). Quantitative analysis revealed a marked difference in mitotic chromosome attachment. Plasmids containing non-peak regions exhibited minimal attachment, with nearly 90% showing less than 50% attachment. In contrast, DNA elements derived from peak regions significantly increased plasmid attachment to mitotic chromosomes, with over 25% displaying at least 50% attachment (Fig. 4f-g). Furthermore, depletion of H3K27ac using A485 also promoted detachment of the engineered plasmids (Extended Data Fig. 8f-g). Notably, the tested peak regions lacked intact coding sequences or expression cassettes, suggesting that plasmid-borne RNAs are unlikely to mediate mitotic interactions, aligning with the finding from RNase A treatment (Extended Data Fig. 7h-m).

Collectively, these data show that ecDNA elements capable of recruiting transcriptional machinery play a functional role in facilitating ecDNA attachment to mitotic chromosomes.

### 7. Mis-segregated ecDNAs are expelled to the cytosol

We further explored the outcomes of mis-segregated ecDNAs. We took advantage of an engineered COLO320DM cell line with a TetO array inserted into *MYC* ecDNAs, which can be visualised with TetR-mNeonGreen using time-lapse live-cell microscopy.^42^ When these cells were treated with dBET6 to degrade BET proteins and induce ecDNA detachment from mitotic chromosomes, mis-segregated ecDNAs were excluded from the primary nuclei in the following cell cycle, with some of which forming micronuclei-like structures in the cytosol (Fig. 5a). This phenomenon was independently validated in SNU16 cells by fixed-cell imaging. H3K27ac depletion by A485 after one cell cycle significantly increased cytosolic DNA FISH signals, which existed in two forms: either dispersed or aggregated in micronuclei (Fig. 5b-c, Extended Data Fig. 9a).

**Fig. 5.**
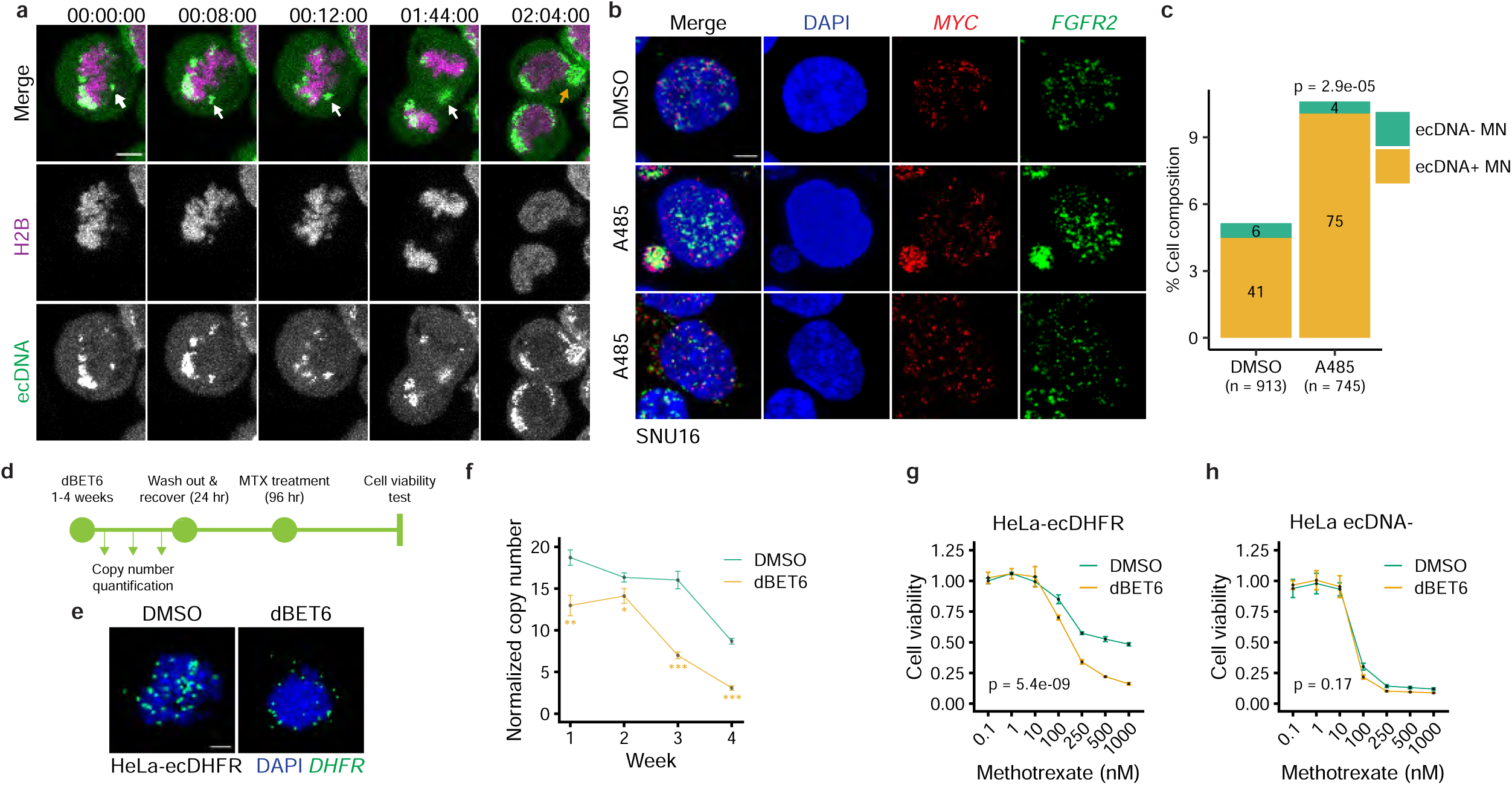
The consequence of mis-segregated ecDNAs. **(a)** Live-cell imaging with COLO320DM TetO-MYC TetR-mNeonGreen cell line under the treatment with 1 μM dBET6 (scale bar: 5 μm. Time scale: 2 hr and 4 min, White arrow: Detached ecDNA. Yellow arrow: Micronuclei-like morphology). **(b)** Representative *MYC* and *FGFR2* ecDNA FISH in SNU16 interphase cells. Cells were treated with 10 μM A485 with a 12-hr nocodazole arrest and were subsequently released to the next cell cycle (scale bar: 5 μm). **(c)** Quantification of micronuclei formation in SNU16 interphase cells (Fisher’s exact test). Cells were treated with 10 μM A485 with a 12-hr nocodazole arrest and were subsequently released for 5 hr to the next cell cycle. Data were pooled from three replicates for statistics. **(d)** Workflow showing the long-term dBET6 treatment for functional assessments. ecDNA copy number was quantified by qPCR every week after treatment. dBET6 was washed out at least 24 hr before cell viability assessment by CCK8 assays. **(e)** Representative *DHFR* ecDNA FISH on HeLa-ecDHFR metaphase cells after 50 nM dBET6 treatment (scale bar: 5 μm). **(f)** qPCR results for *DHFR* ecDNA copy number in HeLa-ecDHFR cells after 1-4 weeks of 50 nM dBET6 treatment (two-sided Student’s t-test). **(g-h)** Methotrexate-cell viability dose-response curves for **(g)** HeLa-ecDHFR and **(h)** parental HeLa cells comparing cells pre-treated with DMSO or dBET6 for 4 weeks (two-way ANOVA).

ecDNA mis-segregation led to its depletion. A long-term A485 treatment in MSTO211H gradually eliminated the number of ecDNAs over 3 weeks and diminished the expression of ecDNA-borne oncogenes, such as *MYC* (Extended Data Fig. 2k & 9b). We then asked whether ecDNA depletion could inhibit cancer aggressiveness, such as therapy resistance. We employed the HeLa-ecDHFR model, in which *DHFR* ecDNAs were induced and maintained by long-term exposure to methotrexate. Degradation of BET proteins by dBET6 also induced *DHFR* ecDNA detachment from mitotic chromosomes (Fig. 5d-e, Extended Data Fig. 9c-e). While the removal of methotrexate decreased *DHFR* ecDNAs in agreement with previous studies,^43^ dBET6 treatment accelerated the loss of *DHFR* ecDNAs and expression (Fig. 5f, Extended Data Fig. 9f). Consequently, ecDNA-depleted HeLa cells were re-sensitised to methotrexate (Fig. 5g). This phenotype was associated with the loss of *DHFR* ecDNAs, rather than the depletion of BET proteins, as parental, ecDNA-free HeLa cells were not further sensitised to methotrexate after long-term exposure to dBET6 (Fig. 5h).

In summary, these data show that mis-segregated ecDNAs were expelled into the cytosol, resulting in the loss of ecDNAs. Depletion of ecDNAs decreases ecDNA-borne oncogene expression and can re-sensitise cancer cells to therapy, whose resistance is driven by ecDNA.

## Discussion

The random segregation of acentric ecDNAs represents a crucial driver of tumour evolution.^16–18^ This process leads to the unequal distribution of ecDNA particles among daughter cells during cell division, thereby fostering tumour heterogeneity. As a result, a diverse pool of cancer cells emerges, each possessing varied oncogene copy numbers that facilitate rapid adaptation to the dynamic tumour microenvironment, particularly under therapeutic pressure.

While previous research has suggested that chromosome hitchhiking mediates ecDNA segregation,^22^ the precise molecular mechanism governing the interaction between ecDNAs and chromosomes during mitosis remains poorly characterised. We propose a model in which ecDNAs preferentially interact with H3K27ac-marked active chromosomal regions during interphase, facilitated by bromodomain proteins, the Mediator complex, and Pol II. As cells enter mitosis, chromosome condensation and transcriptional silencing lead to the eviction of the Mediator complex, while bromodomain proteins and inactive Pol II persist, maintaining ecDNA anchorage to accessible H3K27ac regions (Fig. 3o). This model was validated in cell lines from four cancer types (lung, stomach, colon, and cervix), carrying diverse ecDNA species (*MYC*, *FGFR2*, and *DHFR*), which demonstrates its universality.

Two prior studies have suggested that ecDNAs remain transcriptionally active during mitosis, which is important for ecDNA segregation and co-segregation, as evidenced by the presence of intronic RNA FISH signal and the ability of triptolide to disrupt ecDNA aggregation and chromosome attachment.^19,23^ However, our findings challenge this view. First, ChIP-seq analysis revealed that p-Pol II S2/5, essential for transcription initiation and elongation,^38^ was largely absent from mitotic ecDNAs. Total Pol II was enriched at TSS regions but depleted from gene bodies, indicating stalled transcription (Fig. 3h-i). Second, MED14, an essential structural component of the Mediator complex required for preinitiation complex assembly,^37,44^ was undetectable on mitotic ecDNAs (Fig. 3h-i). Third, 5-EU labelling showed minimal nascent RNA synthesis in ecDNA-containing mitotic cells (Extended Data Fig. 7a). Collectively, these lines of evidence support that ecDNAs are likely transcriptionally inactive during mitosis. This may also explain why post-mitosis-entry depletion of bromodomain protein led to more ecDNA detachment, likely because nascent transcriptional machinery could not be assembled on ecDNAs during mitosis. The observed intronic RNA on mitotic ecDNAs may reflect unspliced, chromatin-associated transcripts synthesised before mitotic entry. Moreover, the effect of triptolide was confounded by its ability to induce proteasome-dependent degradation of Pol II.^45^ Notably, inhibition of transcription elongation using a CDK9 PROTAC, which did not degrade Pol II, failed to detach ecDNAs from mitotic chromosomes (Fig. 3j-n). Therefore, should residual transcriptional activity remain on ecDNAs during mitosis, it is not required for ecDNA tethering to mitotic chromosomes.

Our findings further emphasise that mis-segregated ecDNAs are expelled into the cytosol and gradually depleted, leading to the loss of oncogene expression, thus reversing therapy resistance. This finding opens a new avenue for treating ecDNA-driven cancer by targeting ecDNA segregation. Within the ecDNA segregation machinery, many targets are considered actionable. For example, small molecules targeting histone lysine acetyltransferases, bromodomain proteins, and CDK8 are currently under investigation in clinical trials.^46–49^ Future studies should aim to elucidate whether patients with ecDNA-containing tumours will derive therapeutic benefit from these strategies. Notably, bromodomain proteins exhibit mutually complementary roles, contingent upon the specific cancer model examined. For example, the attachment of *DHFR* ecDNAs in HeLa cells was more sensitive to dBET6 treatment than *MYC* ecDNAs in MSTO211H and SNU16 cells. This raises a critical question regarding whether the segregation of ecDNAs is influenced by specific bromodomain proteins that regulate the expression of their cargo genes.

However, the mechanism mediating cytosolic ecDNAs depletion needed to be further explored. One possibility is nuclease-mediated degradation, such as by TREX1.^50^ Additionally, processes involved in cellular homeostasis, including the autophagy-lysosomal pathway, have been implicated in cytosolic DNA clearance.^51^ Another possibility lies in the replication defect of cytosolic DNAs, which may lead to their dilution through cell division.^50^

An intriguing aspect of our findings is that, even with no perturbations, ecDNA segregation is “leaky” in a few cells, where certain ecDNA particles are not consistently attached to mitotic chromosomes. This phenomenon may lead to the accumulation of cytosolic DNA in ecDNA-driven cancer, which may trigger innate immunity sensing pathways, such as the cGAS/STING pathway.^52^ Although many cancers have evolved mechanisms to silence the expression of cGAS or STING, the relationship between the deactivation of these pathways and ecDNA pathogenesis warrants further investigation. Alternatively, it is also important to understand whether cancer may hijack this pathway for chronic STING activation and promote its growth and metastasis.^1^

One limitation of this study lies in the global depletion approach used for H3K27ac, which hinders the ability to distinguish the specific contributions of mitotic chromosomes and ecDNAs. We partially addressed this by using CRISPRi to target selected ecDNA regions enriched for H3K27ac, as well as BRD4, SMARCA4, and Pol II, achieving significant ecDNA detachment from mitotic chromosomes (Fig. 4a-d). However, current technologies limit simultaneous targeting across all possible loci on ecDNAs. Notably, ecDNA-chromosome mitotic interactions resemble enhancer-promoter contacts, suggesting the functional relevance of the H3K27ac mark on both entities due to its role in enhancer and promoter function.^53^ Another opening question relates to the thermodynamics of ecDNA *trans*-interactions during mitotic entry. As cells progress into prophase, chromatin begins to condense, reducing the number of accessible regions available for ecDNA tethering. The rapid kinetics of enhancer-promoter interactions, occurring within minutes, suggest that ecDNA-chromosome contacts can be efficiently established during early mitosis.^54^ Future studies should explore how ecDNAs dynamically respond to chromatin condensation and spatially reorganise during mitotic progression. Additionally, although long-term depletion of H3K27ac or BET proteins effectively eliminates most ecDNAs, a substantial fraction remains associated with mitotic chromosomes during the first cell cycle of treatment. This suggests the involvement of additional mechanisms in ecDNA segregation. A more comprehensive strategy, such as unbiased profiling and screening as previously reported,^55^ may be necessary to fully elucidate the regulatory landscape governing ecDNA segregation.

## Methods

### Cell culture

MSTO211H, SNU16, COLO320DM, and DLD1 cells were obtained from ATCC. All cells except DLD1 were cultured in Dulbecco’s Modified Eagle’s Medium/F12 (DMEM/F12; Corning) supplemented with 10% Fetal Bovine Serum (FBS; Corning) and 1× Penicillin-Streptomycin (Corning), maintained at 37°C in a humidified incubator with 5% CO_2_, and regularly tested for mycoplasma. DLD1 cells were cultured in RPMI 1640 (Cytiva) supplemented with 10% FBS (Corning) and 1× Penicillin-Streptomycin (Corning), maintained at 37°C in a humidified incubator with 5% CO_2_, and regularly tested for mycoplasma. The HeLa-ecDHFR cell line was a gift from Dr. Roel Verhaak (Yale University), which carries artificially induced *DHFR* ecDNA by long-term exposure to methotrexate. To maintain *DHFR* ecDNAs, 20 μM methotrexate was added to the cell culture media. The COLO320DM TetO-MYC TetR-mNeonGreen cell line was a gift from Dr. Paul Mischel (Stanford University), with its ecDNA inserted with a TetO array and labelled with stably expressed TetR-mNeonGreen.

### Enrichment of metaphase and anaphase cells

Cells were seeded overnight on glass coverslips coated with Poly-D-Lysine (PDL; Sigma-Aldrich). Prophase and prometaphase cells were arrested with 100 ng mL^-1^ nocodazole (Sigma-Aldrich) for 8-12 hr (unless specified) at 37°C. After arrest, cells were washed 3× with pre-warmed PBS and released in growth media for 40-50 min at 37°C, allowing them to progress to metaphase and anaphase. The following compounds were added before or during the nocodazole treatment, whose dosage and timing were indicated in the corresponding main figures: A485 (Selleck Chemicals); dBET6 (Selleck Chemicals or Cayman); AU15330 (Medchemexpress); Actinomycin D (Selleck Chemicals); Triptolide (Selleck Chemicals); α-Amanitin (Santa Cruz Biotechnology); RNase A (New England Biolabs); CDK9i (Medchemexpress).

### Flow cytometry and sorting

Single-cell suspensions were cross-linked with 2% formaldehyde (Sigma-Aldrich) for 10 min, quenched with glycine from respective kits for 10 min, and permeabilised with 0.1% saponin (Thermo Scientific) in PBS for 10 min. After blocking with 1% FBS + 0.1% saponin for 30 min, the anti-phos-MPM2 (Sigma-Aldrich 05-368, 1:1,000) primary antibody was added to the blocking solution and incubated for 1 hr. Cells were then washed 3× with PBS containing 0.1% saponin before AlexaFluor-conjugated secondary antibodies (Invitrogen) staining for 1 hr. After another 3x washes, DNA was stained with 1 μg mL^-1^ DAPI for 5 min. Finally, cells were sorted using the FACSAria Fusion (BD Biosciences) for G1 (2N DNA, MPM2-negative) and M phase (4N DNA, MPM2-positive). All procedures were performed at room temperature (RT). To sort for mitotic ChIP-seq: cells were fixed with 2 mM disuccinimidyl glutarate for 30 min, and then together with 1% formaldehyde (Sigma-Aldrich) for 8 min, and protease inhibitor was added throughout experiment, and F(ab) secondary antibody (Invitrogen) was used to avoid the interference with the subsequent ChIP-seq.

### In situ HiC

*In situ* HiC libraries for G1 and M phases sorted cells were prepared with the Arima HiC kit (Arima Genomics) following the manufacturer’s instructions. In brief, one million sorted cells, which were cross-linked with 2% formaldehyde before sorting, were digested with two restriction enzymes, followed by end filling, biotin labeling, and proximal ligation. Next, 1.5 μg ligated DNA was fragmentised by the Covaris ME220 (peak power: 75W; duty factor, 20%; cycles per burst: 200; treatment time, 3.5 min) for 300-bp size selection with DNA purification beads. Biotin-labeled DNA was then enriched with streptavidin beads for sequencing library preparation using the Arima library prep module. HiC libraries were then pooled and sequenced with the paired-end 150 mode.

Adapter sequences were trimmed by the fastp software (v0.22.0). Trimmed reads were then processed through the HiC-Pro pipeline (v3.1.0) with hg38 as the reference genome. The reproducibility of HiC libraries was analysed by the HiCRep software (v0.2.6). HiC interaction was detected by the FitHiC2 software (v2.0.8).

### CUT&RUN sequencing

CUT&RUN libraries were generated with the CUTANA ChIC/CUT&RUN kit version 4 (EpiCypher, 12-1048) and the CUT&RUN library prep kit (EpiCypher, 14-1001&14-1002) according to their instructions. MSTO211H cells were cultured and treated as described above in 10-cm plates. Cells were harvested by trypsinisation and counted with Countess 3 Automated Cell Counter (Invitrogen). For one CUT&RUN sample preparation, five hundred thousand cells were used to isolate nuclei using the CUTANA Pre-Nuclei Extraction Buffer (EpiCypher, 21-1026a). Nuclei were then bound to beads and incubated with 0.5 µg of antibodies overnight. The following antibodies were used: IgG (EpiCypher 13-0042), and anti-H3K27ac (EpiCypher 13-0059). Target DNAs bound by antibody were digested with pAG-MNase for 2 hr at 4°C and extracted by incubating with stop master mix containing 0.5 ng of *Escherichia coli* Spike-in DNA at 37°C.

CUT&RUN sequencing data were analysed as previously described. In brief, sequencing data were aligned to the hg38 human genome. Unmapped and discordant read pairs were removed. The coverage was normalised with the spike-in control, which was determined by the abundance of reads aligned to the *Escherichia coli* genome.

### Chromatin immunoprecipitation sequencing (ChIP-seq)

ChIP-seq libraries were prepared using the True MicroChIP-seq Kit (Diagenode, for flow cytometer-sorted mitotic cells) and SimpleChip Plus Enzymatic Chromatin IP Kit (Cell Signaling Technology, for BRD4 and SMARCA4 ChIP where PROTAC inhibitors were applied), following the manufacturer’s instructions. The following antibodies were used: anti-H3K4me3 (Diagenode C15410003), anti-H3K27ac (Epicypher 13-0059), anti-BRD4 (Bethyl Laboratories A301-985A), anti-SMARCA4 (Cell Signaling Technology 49360S for mitotic cells, Abcam 110641 for interphase cells), anti-MED14 (Abcam ab72141), anti-RPB1 (Cell Signaling Technology 2629S), anti-phospho-RPB1 Ser2/5 (Cell Signaling Technology 13546S). For BRD4 and SMARCA4 ChIP where PROTAC inhibitors were applied, spike-in chromatin and antibody (Active Motif) were added to the reaction. ChIP-seq data were aligned to the human hg38. For BRD4 and SMARCA4 ChIP-seq with PROTAC treatments, data were aligned to the dm6 Drosophila reference genome for spike-in normalisation. ecDNA signals were normalised with copy number by the bigwigCompare function (--operation ratio) from deepTools (v3.5.5) software, with the ChIP-seq input serving as a surrogate for whole-genome coverage.

### RNA interference

Forty nanomolar of respective siRNAs were mixed with Lipofectamine RNAiMAX Transfection Reagent (Invitrogen) following the manufacturer’s instructions and incubated for 10 min at RT to form siRNA-lipid complexes. siRNA-lipid complexes were added dropwise into the cell culture, and gene knockdown was determined 48 hr after transfection. Information about siRNAs is provided in Supplementary Table S1.

### Immunofluorescence

Cells were rinsed with PBS and fixed using 4% PFA for 20 min at RT. Fixed samples were permeabilised with 0.5% Triton X-100 in PBS for 20 min at RT. The following primary antibody was diluted in Renaissance background reducing diluent (Biocare Medical) and incubated overnight at 4°C: anti-H3S10phS28ph 1:400 (Active Motif, 39148). Cells were then washed 3x with PBS for 5 min. Alexa fluor-conjugated secondary antibodies (Invitrogen) were diluted 1:1000 in the same diluent and incubated for 1 hr at RT. After 3x PBS wash, DNA was counterstained with1 μg mL^-1^ DAPI for 5 min. Coverslips were mounted with VECTASHIELD Antifade Mounting Medium (Vector Laboratories) and sealed for imaging. All procedures were performed at RT unless indicated otherwise.

### DNA FISH

Cells on coverslips were rinsed with PBS and fixed using fresh Carnoy’s fixative (3:1 methanol:acetic acid) for 10 min. Fixed cells were air-dried for 2 hr at RT or overnight at 4°C. Samples were equilibrated with 2× SSC and dehydrated in 70%, 85% and 100% ethanol for 2 min each. FISH probes (Empire Genomics) diluted in hybridisation buffer (1:10) were applied to coverslips and mounted onto glass slides. Slides were denatured for 2 min at 75°C and hybridised overnight at 37°C in a humidified chamber. After removing from slides, coverslips were washed 2× in 0.4× SSC with 0.3% IGEPAL for 5 min, followed by another wash in 2× SSC with 0.1% IGEPAL for 5 min. DNA was counterstained with 1 μg mL DAPI in 2× SSC for 5 min and washed 2× with 2× SSC for 5 min. Coverslips were mounted onto glass slides with VECTASHIELD antifade mounting medium (Vector Laboratories) and sealed for imaging. All procedures were performed at RT unless indicated otherwise.

### DNA Oligopaint

Cells on coverslips were rinsed with PBS and fixed using fresh Carnoy’s fixative (3:1 methanol:acetic acid) for 10 min. Fixed cells were air-dried for 2 hr at RT or overnight at 4°C. Samples were equilibrated with 2× SSC and dehydrated in 70%, 85% and 100% ethanol for 2 min each. Primary oligopaint probes (Designed in this study, Twist Bioscience) diluted in hybridisation buffer (1:10) were applied to coverslips and mounted onto glass slides. Slides were denatured for 2 min at 75°C and hybridised overnight at 37°C in a humidified chamber. After removing from slides, coverslips were washed 2× in 0.4× SSC with 0.3% IGEPAL for 10 min at 45°C, followed by another wash in 2× SSC with 0.1% IGEPAL for 10 min at RT. Secondary oligopaint probes (Designed in this study, Sigma-Aldrich) diluted in hybridisation buffer (1:10) were applied to coverslips and mounted onto parafilm. The coverslips were hybridised for 2 hr at 37°C in a humidified chamber. After removing from slides, coverslips were washed 2× in 0.4× SSC with 0.3% IGEPAL for 10 min at 45°C, followed by another wash in 2× SSC with 0.1% IGEPAL for 10 min at RT. DNA was counterstained with 1 μg mL^-1^ DAPI in 2× SSC for 5 min and washed 2× with 2× SSC for 5 min. Coverslips were mounted onto glass slides with VECTASHIELD antifade mounting medium (Vector Laboratories) and sealed for imaging. All procedures were performed at RT unless indicated otherwise.

### Immunofluorescence and DNA Oligopaint

Immunofluorescence was performed first, following the immunofluorescence section above. The following primary antibody was used: anti-H3K27ac 1:50 (Cell Signaling Technology 8173S). After secondary antibody staining, the cells were fixed with 4% formaldehyde in PBS for 15 min at RT. Fixed cells were dehydrated using fresh Carnoy’s fixative (3:1 methanol:acetic acid) for 15 min at RT. Samples were rinsed with 100% ethanol and were air-dried at RT. Primary oligopaint probes (Designed in this study, Twist Bioscience) diluted in hybridisation buffer (1:10) were applied to coverslips and mounted onto glass slides. Slides were denatured for 20 min at 75°C and hybridised overnight at 37°C in a humidified chamber. The rest of the procedure were the same as the DNA oligopaint section above.

### Cytospin and immunofluorescence

Cells were arrested in metaphase with 33 ng mL^-1^ KaryoMAX colcemid (Gibco) overnight at 37°C and pelleted in hypotonic buffer (10 mM Tris-HCl (pH 7.5) + 20 mM NaCl + 5 mM KCl + 1 mM CaCl_2_ + 0.5 mM MgCl_2_ + 40 mM glycerol) for 15 min on ice. 0.1% Tween-20 was then added to the hypotonic buffer to permeabilize cells. Cells were deposited onto glass slides using Cytospin 4 Cytocentrifuge (Epredia) at 500 RPM for 10 min. Slides were aged overnight at 4°C and blocked with Renaissance background reducing diluent (Biocare Medical) for 10 min at RT. The following primary antibodies were diluted in the diluent and incubated for 1 hr at RT: anti-H3K27ac 1:100 (Cell Signaling Technology 8173S) and anti-H3K4me3 1:100 (Cell Signaling Technology 9751S). Slides were washed 3× with the diluent containing 0.1% Tween-20 for 5 min before AlexaFluor-conjugated secondary antibody staining (Invitrogen) for 1 hr at RT. Following another 3× washes as above, 1 μg mL^-1^ DAPI in the diluent was applied for DNA counterstaining. Cells were then fixed with 4% formaldehyde for 15 min at RT before being mounted with VECTASHIELD antifade mounting medium and sealed for imaging.

### RNA abundance detection

Cells on coverslips were treated with 1 mg mL^-1^ RNase A for 40 min. Total RNA abundance was detected with Cell Navigator Live Cell RNA Imaging Kit (AAT Bioquest) following the manufacturer’s instructions. In brief, cells were fixed with methanol and permeabilised with 1% Triton X-100. After washing with PBS, the samples were incubated with RNA dye according to the instructions and stained with 1 μg mL^-1^ DAPI. The slides were then mounted and imaged. The RNA signal intensity was quantified with ImageJ.

### Nascent RNA detection

Cells were treated with nocodazole for 7 hr. Then the medium was replaced with medium containing 1 mM of 5-EU (Vector Laboratories) for 1 hr at 37°C. The cells were then fixed using fresh Carnoy’s fixative (3:1 methanol:acetic acid) for 10 min. After DNA FISH, the cells were treated with a click reaction using Click-&-Go Plus 488 assay kit (Vector Laboratories), followed by DAPI staining and mounting.

### Fluorescence microscopy

DNA FISH and immunofluorescence were imaged on the Zeiss Axio Observer 7 microscope equipped with the Apotome 3 optical sectioning module. Images were acquired with a 63× Plan-Apochromat 1.40 NA oil objective and processed using ZEN software (v3.4, Zeiss). At least five z-stacks were taken with an interval of 1 μm. Maximum projection was applied to generate a 2D representation of the assay.

For live-cell imaging, COLO320DM TetO-MYC TetR-mNeonGreen cells were seeded in 35-mm glass-bottom dishes and treated with 1000 nM dBET6. Culture media were replaced with warm FluoroBrite DMEM (Gibco) before imaging. Images were acquired with the Zeiss LSM 980 microscope through a 63× oil objective. Three slices per 1-μm interval were recorded every 4 min.

### Immunoblotting

Cells were lysed in RIPA buffer supplemented with 1× protease inhibitor cocktail (Sigma-Aldrich) and 1× phosphatase inhibitor cocktail (Sigma-Aldrich). To improve histone extraction efficiency, lysates were added with 100 units μL^-1^ of Micrococcal Nuclease (New England Biolabs) and 1 mM of calcium chloride and incubated for 15 min at RT. Lysates were then cleared by centrifugation (17000 RCF for 15 min at 4°C) and quantified using Pierce Rapid Gold BCA Protein Assay Kit (Thermo Scientific). Samples were resolved by SDS polyacrylamide gel electrophoresis and transferred onto 0.2-μm nitrocellulose membranes (Bio-Rad). Membranes were blocked with 2% Bovine Serum Albumin (BSA) in TBST (0.1% Tween-20 in TBS) for 1 hr at RT. The following primary antibodies were diluted in 2% BSA in TBST and incubated with membranes overnight at 4°C: anti-BRD2 1:1,000 (Bethyl Laboratories A302-583A), anti-BRD3 1:1,000 (Bethyl Laboratories A700-069), anti-BRD4 1:1,000 (Bethyl Laboratories A301-985A), anti-SMARCA4 1:1,000 (Cell Signaling Technology 49360S), anti-H3K27ac 1:1,000 (Cell Signaling Technology 8173S), anti-H3K4me3 1:1,000 (Cell Signaling Technology 9751S), anti-RPB1 1:1,000 (Cell Signaling Technology 2629S), anti-phospho-RPB1 Ser2/5 1:1,000 (Cell Signaling Technology 13546S), anti-c-MYC 1:2,000 (Abcam ab32072), anti-CDK9 1:1,000 (Cell Signaling Technology 2316S), anti-DHFR 1:1,000 (Cell Signaling Technology 43497), anti-Vinculin 1:10,000 (Proteintech 66305-1-Ig), and anti-GAPDH 1:10,000 (Proteintech 60004-1-Ig). After primary antibody incubation, membranes were washed 3× with TBST and incubated with HRP-conjugated secondary antibodies (Cell Signaling Technology) diluted to 1:5,000 with 2% BSA in TBST for 1 hr at RT. Membranes were washed 3× with TBST, developed using SuperSignal West Pico PLUS Chemiluminescent Substrate (Thermo Scientific), and imaged using ImageQuant 800 (Cytiva Amersham).

### qPCR

For mRNA reverse transcription qPCR, RNA was extracted using the Quick-RNA Miniprep Plus Kit (Zymo Research) and quantified by NanoDrop (Thermo Scientific). One microgram of RNA was reverse transcribed to cDNA using the Maxima H Minus reverse transcriptase (Thermo Scientific). Ten nanogram cDNA was added with 1× SYBR Green qPCR Master Mix (Selleck Chemicals) and 400 nM respective forward and reverse primers for qPCR on CFX Opus 96 System (Bio-Rad). The relative RNA level was calculated by the ddCt method. The *TBP* transcript was used as a control for most assays, except for the transcription inhibitor assessment, in which the long half-life *RPL13A* transcript was used. Primer sequences are listed in Supplementary Table S1.

For DNA copy number qPCR, genomic DNA was extracted using the Quick-DNA Miniprep Plus Kit (Zymo Research) and quantified by the Qubit assay (Thermo Scientific). Ten nanograms of DNA was added with 1× SYBR Green qPCR Master Mix (Selleck Chemicals) and 400 nM respective forward and reverse primers for qPCR on CFX Opus 96 System. The gene copy number was determined using *HBB* as a diploid control.

### Plasmid cloning and transfection

Engineered plasmids were made based on pSP2-96-merTetO-EFS-BLaR (Addgene 118713). The parental plasmids were cut with BamHI-HF (New England Biolabs) and NdeI (New England Biolabs) restriction enzymes and purified as backbones. The ecDNA regions were obtained through PCR amplification using MSTO211H genomic DNA and were cloned into the backbone using NEBuilder HiFi DNA Assembly Kit (New England Biolabs). DLD1 cells were seeded overnight in a 6-well plate. Two micrograms of plasmids were transfected into DLD1 cells with X-tremeGENE HP DNA Transfection Reagents (Roche). Twelve hr post-transfection, the cells were passaged on glass coverslips in a 6-well plate. Twenty-four hr later, prophase and prometaphase cells were arrested with 100 ng mL^-1^ nocodazole (Sigma-Aldrich) for 8 hr at 37°C. After arrest, cells were washed 3× with pre-warmed PBS and released in growth media for 40 min at 37°C, allowing them to progress to metaphase and anaphase. Then the cells were fixed and used for DNA FISH.

### Cell viability assay

Five thousand cells/well HeLa-ecDHFR or HeLa parental cells were seeded into 96-well plates. Cells were treated with methotrexate for 4 days. Ten microliter CCK-8 reagents (Apexbio Technology) were added and incubated at 37°C for 2 hr. Cell viability was determined as the absorbance at 450 nm using a plate reader.

### CRISPR interference

The dCas9 expression vector (Addgene 71236) was packaged into lentivirus. Then, MSTO211H cells were infected with lentivirus to stably express dCas9 proteins. The sgRNA vectors were constructed using LentiGuide-Puro (Addgene 52963) as a backbone and then packaged into lentivirus. In each experiment, the MSTO211H-dCas9 cell line was infected with sgRNA-expression virus upon seeding, together with 10 μg mL^-1^ polybrene transfection reagent (Sigma-Aldrich). Twenty-four hr post-infection, the medium is changed to remove the excess virus. Forty-eight hr post-infection, the cells were seeded onto coverslips coated with PDL. Seventy-two hr post-infection, nocodazole was added to arrest the cell for 8 hr. Then the cells were fixed, and DNA FISH was performed to check ecDNA in mitotic cells.

### Statistics

Experiments were repeated at least three times. All statistical methods are indicated in the corresponding figure legends. Student’s t-test, one-way ANOVA, or two-way ANOVA was applied for normally distributed, homoscedastic data. Otherwise, the Wilcoxon or Kruskal-Wallis test was used. All statistical tests were two-sided. P values from multi-comparison tests were adjusted using the false discovery rate (FDR) method. Bar plots are shown as mean ± SD. Box plots are shown as median and upper/lower quartiles, with whiskers showing the 1.5× interquartile range. In the ECDF plot, dAUC was shown as mean ± SEM, and p values were calculated based on the pooled data. Each point in the dot plot represents a single cell or a replicate. Different shapes of dots were used to indicate different replicates in the dot plot. A density plot displaying the data distribution of each group is shown when applicable.

## Data availability

HiC, CUT&RUN, and ChIP-seq data are deposited into the NIH Sequence Read Archive under BioProject accession PRJNA1263546.

## Acknowledgments

**(a)** S. Wu is supported by the Cancer Prevention and Research Institute of Texas (CPRIT, RR210034), the American Cancer Society (CAT-24-1379043-01-CAT), and the Gilead’s Research Scholars Program in Solid Tumors. J.Y.S. Lim. is supported by the CPRIT Training Grant (RP210041). This work was delivered as part of the eDyNAmiC team supported by the Cancer Grand Challenges partnership funded by Cancer Research UK (P.S.M CGCATF-2021/100012, S.W. CGCATF-2021/100023) and the National Cancer Institute (P.S.M. OT2CA278688, S.W. OT2CA278683). Certain graphic elements were obtained from the open-sourced Bioicons (CC-BY 3.0 unported license). We thank Dr. Pippa Cosper at the University of Wisconsin for her advice on this project.

## Author contributions

**(a)** Y. Xie and S. Wu conceptualised the research. Y. Xie, J.Y.S. Lim, and S. Wu developed the methodology and analysed data. Y. Xie, J.Y.S. Lim, W. Liu, C. Gilbreath, X. Sun, and Y.J. Kim performed experiments. K. Qiao helped with figure design. Y. Xie and S. Wu wrote the manuscript. Y.J. Kim and S. Wu supervised this research.

## Extended data figure legends

**Extended Data Fig. 1.**
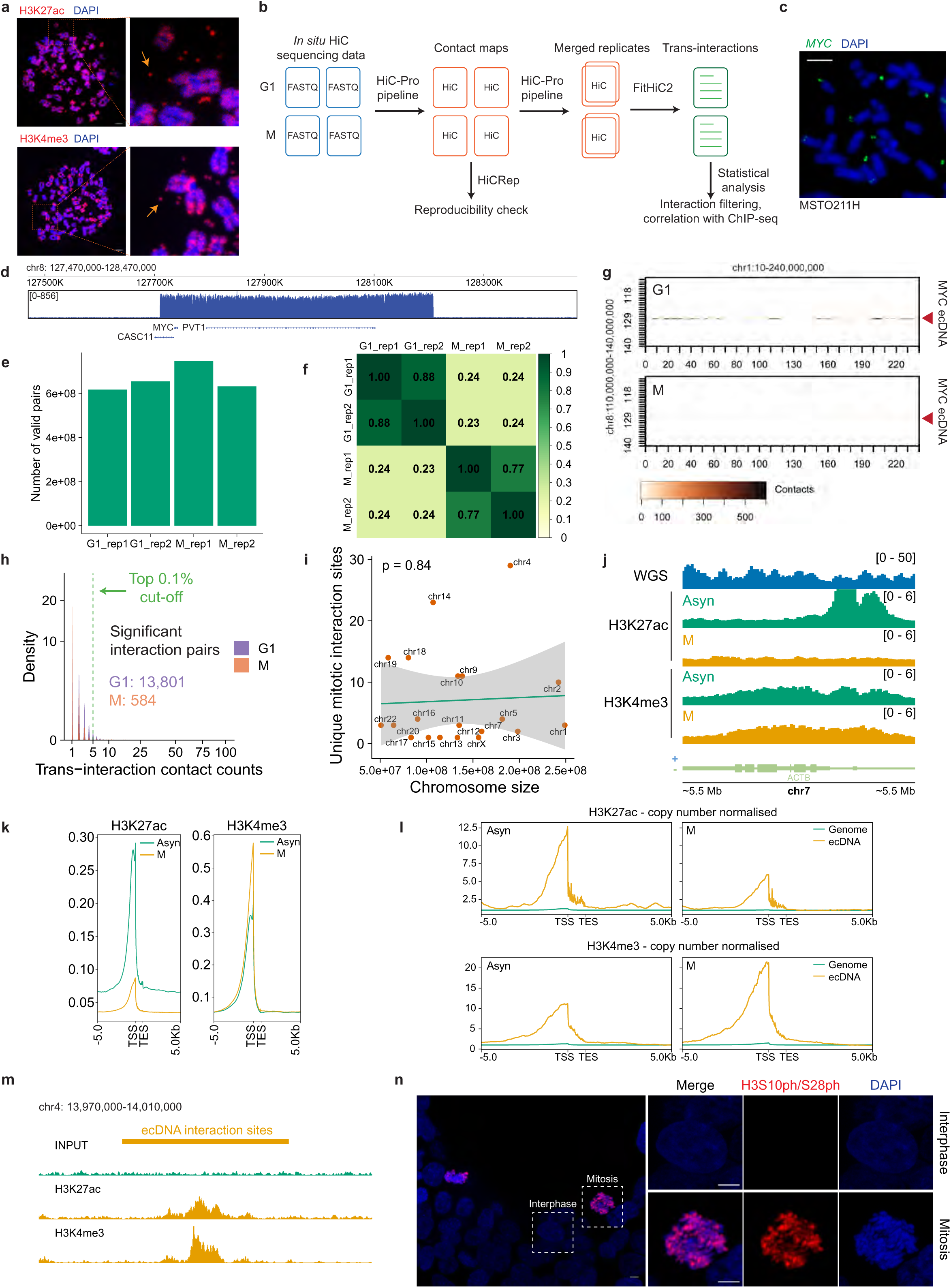
HiC analysis to identify the epigenetic features of ecDNA *trans*-interaction sites during mitosis. **(a)** Abundant H3K27ac and H3K4me3 are present on ecDNAs at metaphase, determined by immunofluorescence of MSTO211H cells. Arrows indicate ecDNAs (scale bar: 5 μm). **(b)** Workflow of integrative analysis of HiC and ChIP-seq data. Two HiC replicates from G1 and M phases sorted cell fraction were processed by the HiC-Pro pipeline to obtain contact maps. After the reproducibility check, replicates from the same cell cycle fraction were merged for *trans*-interaction analysis. Top interaction loci were extracted for subsequent ChIP-seq coverage analysis. **(c)** Representative *MYC* ecDNA FISH image on MSTO211H cells at metaphase showing *MYC* ecDNAs (scale bar: 5 μm). **(d)** WGS track of MSTO211H cells showing the *MYC* ecDNA locus. **(e)** Number of valid pairs of each HiC library. **(f)** Stratum-adjusted correlation coefficient matrix among HiC libraries. **(g)** HiC contact heatmap showing decreased *MYC* ecDNA *trans*-interactions with chromosome 1 in the M phase (bottom panel) compared to the G1 phase (top panel). **(h)** Density distribution of ecDNA *trans*-interaction contact counts in G1 and M phases in MSTO211H cells. The top 0.1% ecDNA-derived *trans*-interactions were defined as significant interaction sites. **(i)** Distribution of ecDNA *trans*-interaction sites on different chromosomes. **(j)** ChIP-seq coverage tracks for H3K27ac, and H3K4me3 at *ACTB* locus in asynchronous (Asyn) or mitotic (M) MSTO211H cells. **(k)** H3K27ac and H3K4me3 ChIP-seq coverage profiles in asynchronous (Asyn) and mitotic (M) MSTO211H cells. TSS: Transcription start site; TES: Transcription end site. **(l)** Quantification for H3K27ac, and H3K4me3 ChIP-seq read counts on ecDNA versus the chromosomal genes in asynchronous (Asyn) or mitotic (M) MSTO211H cells. **(m)** Representative H3K27ac and H3K4me3 ChIP-seq coverage tracks of the ecDNA interaction sites in mitotic MSTO211H cells. **(n)** Representative immunofluorescence images for H3S10ph/S28ph in interphase and mitotic MSTO211H cells (scale bar: 5 μm).

**Extended Data Fig. 2.**
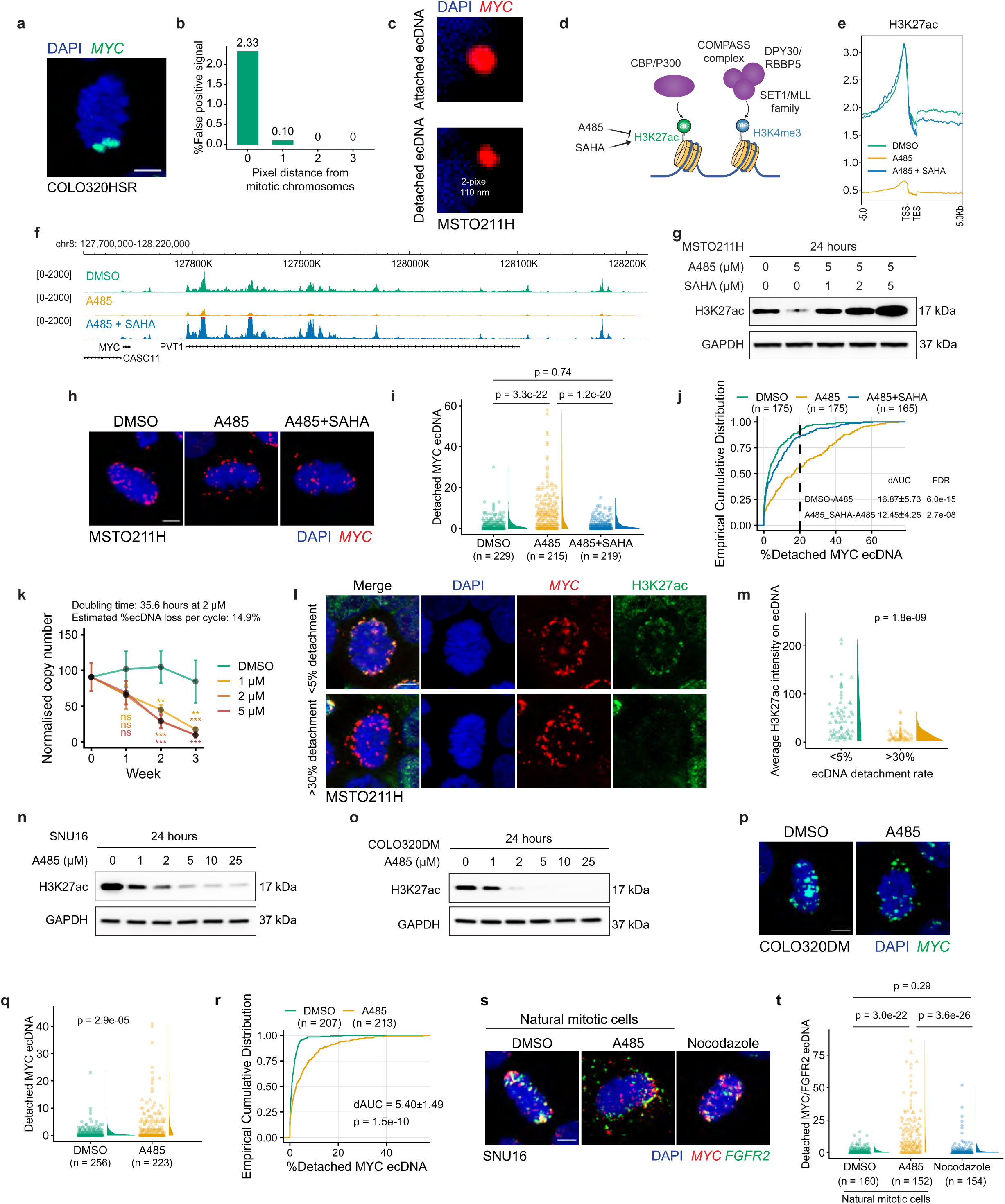
H3K27ac is required for ecDNA attachment to mitotic chromosomes. **(a)** Representative *MYC* ecDNA FISH image on COLO320HSR cells at metaphase showing *MYC* ecDNAs (scale bar: 5 μm). **(b)** The percentage of false positive signals when the pixel distance from the chromosomes is set as 0-3. **(c)** Representative *MYC* ecDNA FISH images on MSTO211H cells at metaphase to show the 2-pixel distance between detached ecDNA and chromosomal DNA. **(d)** Illustration depicting the writers of H3K27ac and H3K4me3 histone modifications. **(e)** H3K27ac CUT&RUN coverage profiles in DMSO, 5 μM A485, and 5 μM A485+2 μM SAHA-treated MSTO211H cells. TSS: Transcription start site; TES: Transcription end site. **(f)** Representative CUT&RUN coverage tracks on the ecDNA region for H3K27ac after 5 μM A485 and 2 μM SAHA treatment in MSTO211H cells. **(g)** Western blot for H3K27ac in MSTO211H cells after A485 and SAHA treatment. **(h)** Representative *MYC* ecDNA FISH on 5 μM A485 and 2 μM SAHA treated MSTO211H metaphase cells (scale bar: 5 μm). **(i-j)** Increased number **(i)** and percentage **(j)** of detached *MYC* ecDNAs per MSTO211H metaphase cell after 5 μM A485 treatment and rescued by 2 μM SAHA treatment (Kruskal-Wallis test with post-hoc Dunn test). The vertical dashed line indicates the percentage of cells with more than 20% detached ecDNA. Nocodazole treatment: 8-12 hr. Data were pooled from three replicates for statistics. dAUC was shown as mean ± SEM. **(k)** qPCR results showing continuously decreased *MYC* ecDNA copy number in MSTO211H cells after 1-3 weeks of A485 treatment (one-way ANOVA with post-hoc Dunnett test). **(l)** Representative *MYC* ecDNA FISH and H3K27ac immunofluorescence on 5 μM A485 treated MSTO211H metaphase cells (scale bar: 5 μm). **(m)** Relative H3K27ac intensity on *MYC* ecDNA in 5 μM A485-treated metaphase MSTO211H cells. Quantification was performed for the cells with high versus low ecDNA detachment rates (two-sided Wilcoxon test). Nocodazole treatment: 8 hr. Data were pooled from two replicates for statistics. **(n)** Western blot showing a dose-response of H3K27ac downregulation after A485 treatment in SNU16 cells. **(o)** Dose-responding H3K27ac downregulation after A485 treatment in COLO320DM cells measured with Western blot. **(p)** Representative *MYC* ecDNA FISH on 10 μM A485-treated COLO320DM metaphase cells (scale bar: 5 μm). **(q-r)** The number **(q)** and percentage **(r)** of detached *MYC* ecDNAs per COLO320DM metaphase cell after 10 μM A485 treatment (two-sided Wilcoxon test). Nocodazole treatment: 4-6 hr. Data were pooled from three replicates for statistics. dAUC was shown as mean ± SEM. **(s)** Representative *MYC* and *FGFR2* ecDNA FISH on 10 μM A485-treated natural SNU16 metaphase cells (scale bar: 5 μm). **(t)** The number of detached *MYC* and *FGFR2* ecDNA per natural SNU16 metaphase cell after A485 treatment (Kruskal-Wallis test with post-hoc Dunn test). Nocodazole treatment: 12 hr. Data were pooled from two replicates for statistics.

**Extended Data Fig. 3.**
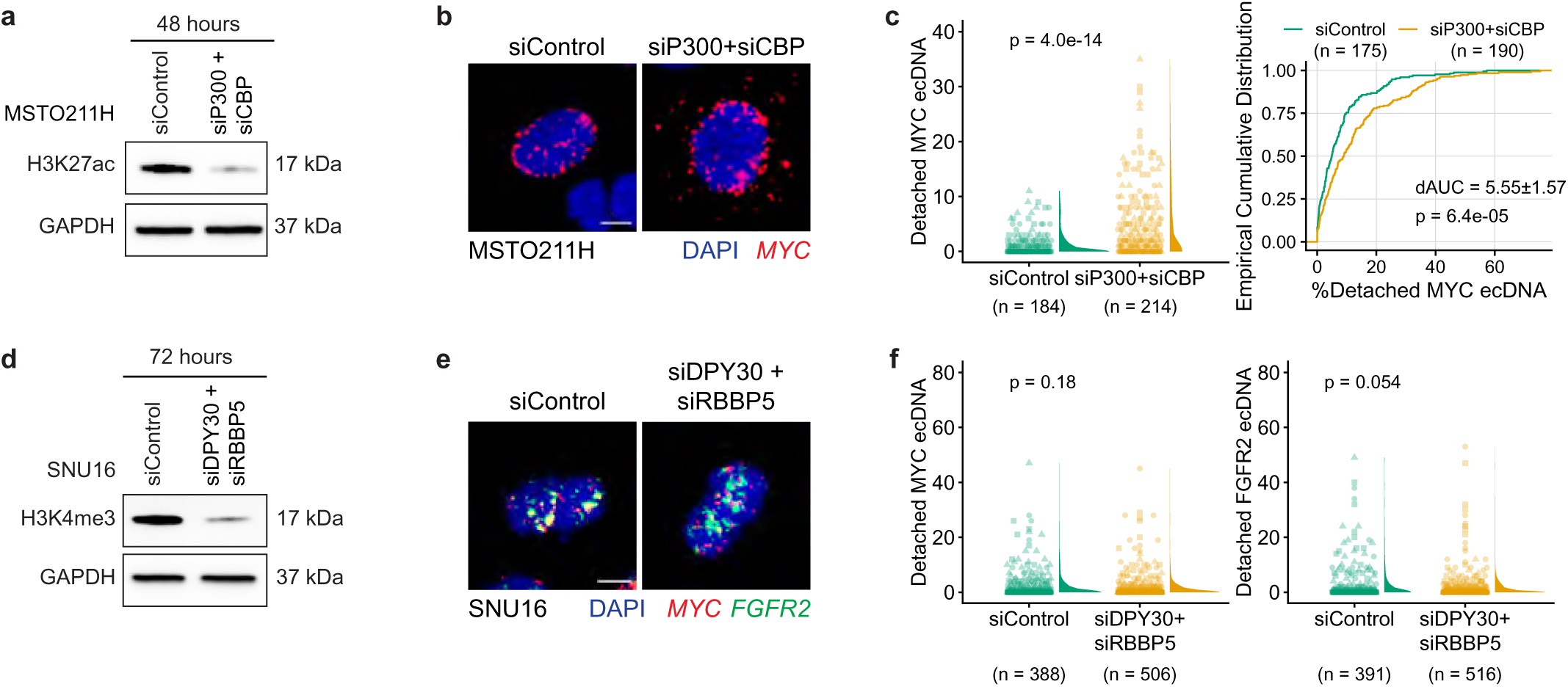
H3K4me3 is not required for ecDNA attachment to mitotic chromosomes. **(a)** Western blot showing decreased H3K27ac abundance in MSTO211H cells after *CBP* and *P300* RNAi knockdown. **(b)** Representative *MYC* ecDNA FISH on MSTO211H metaphase cells after *CBP* and *P300* knockdown (scale bar: 5 μm). **(c)** The number and percentage of detached *MYC* ecDNAs per MSTO211H metaphase cell after *CBP* and *P300* knockdown (two-sided Wilcoxon test). Nocodazole treatment: 8-12 hr. Data were pooled from three replicates for statistics. dAUC was shown as mean ± SEM. **(d)** Western blot showing decreased H3K4me3 after *DPY30* and *RBBP5* RNAi knockdown in SNU16 cells. **(e)** Representative *MYC* and *FGFR2* ecDNA FISH on SNU16 metaphase cells after *DPY30* and *RBBP5* knockdown (scale bar: 5 μm). **(f)** The number of detached *MYC* and *FGFR2* ecDNA per SNU16 metaphase cell after *DPY30* and *RBBP5* RNAi knockdown (two-sided Wilcoxon test). Nocodazole treatment: 12-16 hr. Data were pooled from three replicates for statistics.

**Extended Data Fig. 4.**
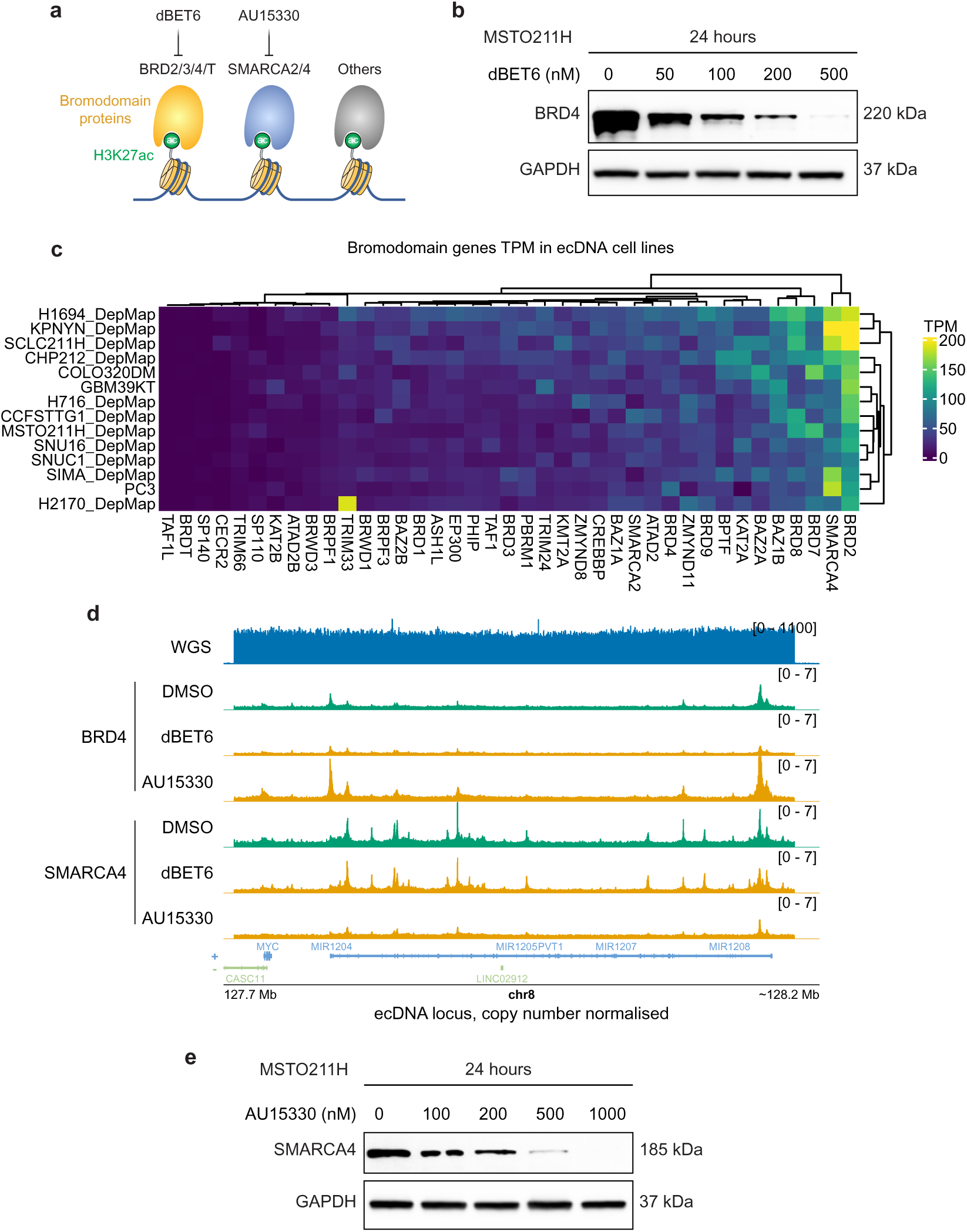
Manipulation of the bromodomain proteins. **(a)** Illustration of major bromodomain-containing proteins as the readers of H3K27ac histone modification. **(b)** Western blot showing dose-responding degradation of BRD4 after dBET6 treatment in MSTO211H cells. **(c)** Heatmap for the mRNA expression level of all human bromodomain-containing proteins in ecDNA-containing cancer cell lines. Data were obtained from public RNA-seq. TPM: transcripts per million. **(d)** Representative ChIP-seq coverage tracks for BRD4 and SMARCA4 after 200 nM dBET6 and 1 μM AU15330 treatment in MSTO211H cells. Values are log2 normalised over corresponding inputs. **(e)** Western blot showing dose-responding degradation of SMARCA4 after AU15330 treatment in MSTO211H cells.

**Extended Data Fig. 5.**
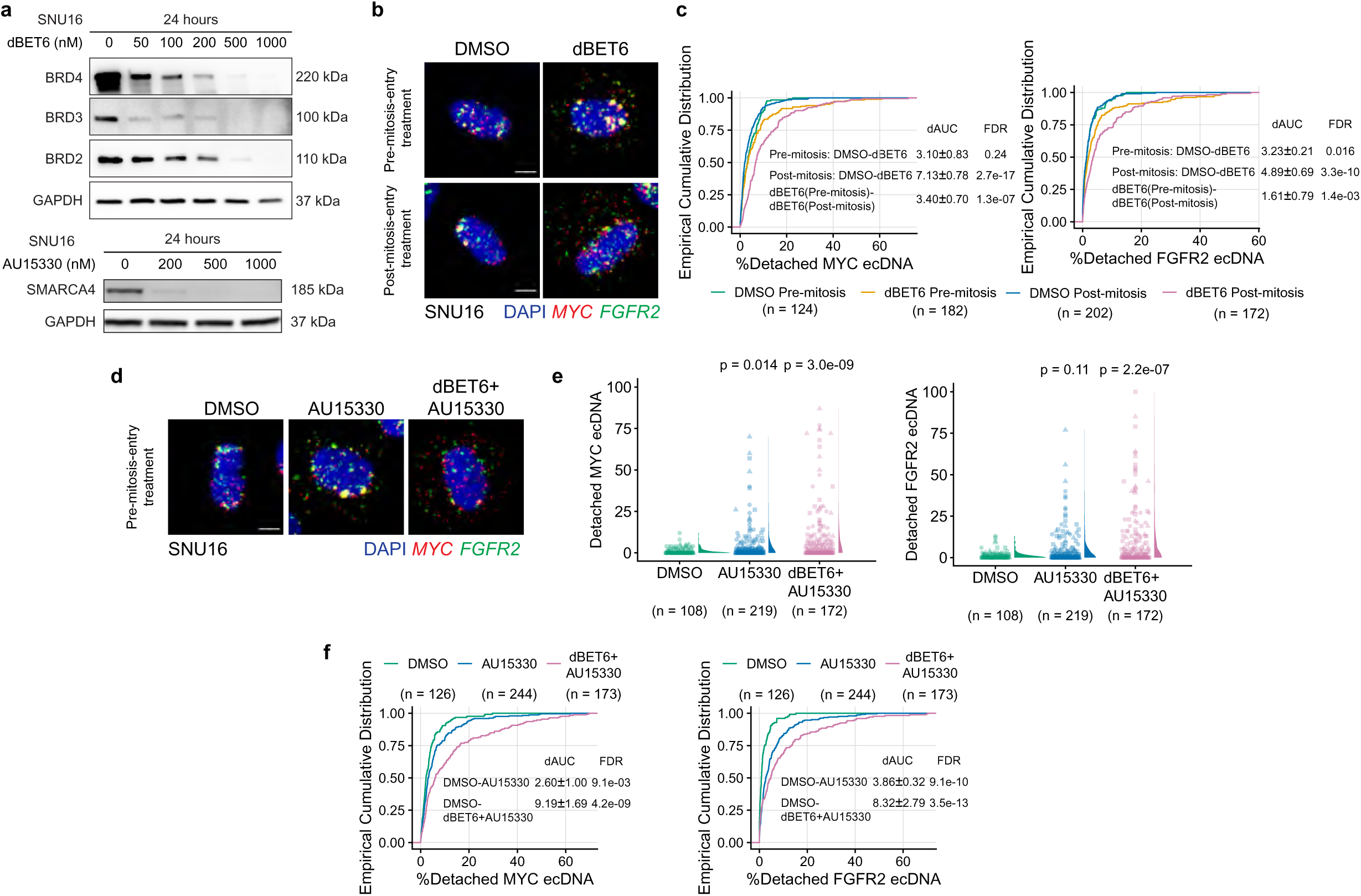
Bromodomain proteins are functionally redundant for ecDNA attachment. **(a)** Western blot for BRD2, BRD3, and BRD4 in SNU16 cells after dBET6 treatment for 24 hr or SMARCA4 after AU15330 treatment for 24 hr. **(b)** Representative *MYC* and *FGFR2* ecDNA FISH on 100 nM dBET6 (pre-mitosis-entry) or 1 μM dBET6 (post-mitosis-entry) treated SNU16 metaphase cells (scale bar: 5 μm). **(c)** The percentage of detached *MYC* and *FGFR2* ecDNAs per SNU16 metaphase cell after pre-mitosis-entry or post-mitosis-entry dBET6 treatment (Kruskal-Wallis test with post-hoc Dunn test). Nocodazole treatment: 8 hr. Data were pooled from three replicates for statistics. dAUC was shown as mean ± SEM. **(d)** Representative *MYC* and *FGFR2* ecDNA FISH on 1 μM AU15330 or 100 nM dBET6+1 μM AU15330 treated (pre-mitosis-entry) SNU16 metaphase cells (scale bar: 5 μm). **(e-f)** The number **(e)** and percentage **(f)** of detached *MYC* and *FGFR2* ecDNAs per SNU16 metaphase cell after AU15330 or dBET6+AU15330 pre-mitosis-entry treatment (Kruskal-Wallis test with post-hoc Dunn test). Nocodazole treatment: 8 hr. Data were pooled from three replicates for statistics. dAUC was shown as mean ± SEM.

**Extended Data Fig. 6.**
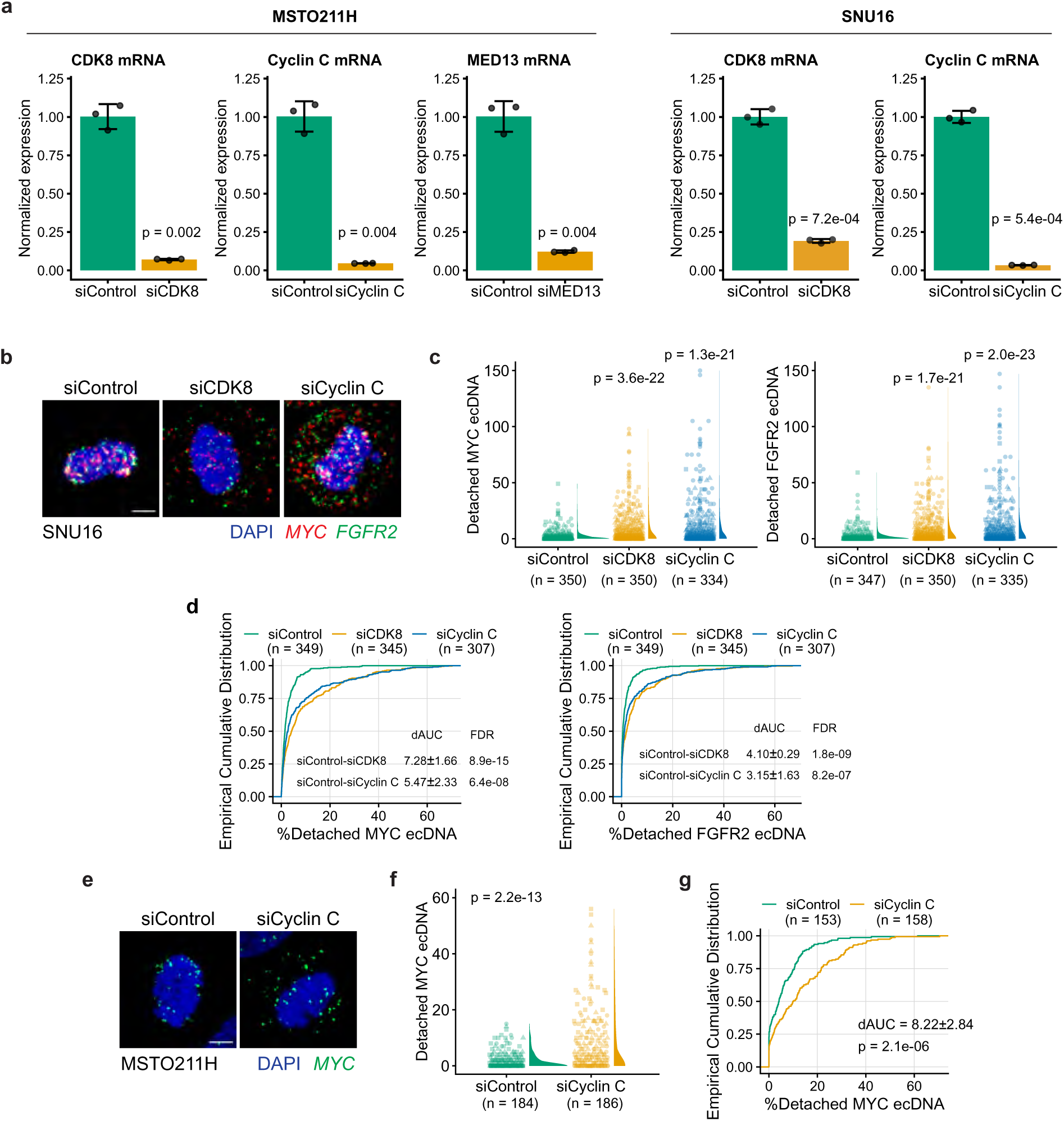
The kinase module of the Mediator complex regulates ecDNA segregation. **(a)** *CDK8*, Cyclin C, and *MED13* RNAi knockdown efficiency assessed by qPCR in MSTO211H and SNU16 cells (two-sided Student’s t-test). **(b)** Representative *MYC* and *FGFR2* ecDNA FISH in SNU16 metaphase cells after Cyclin C or *CDK8* RNAi knockdown (scale bar: 5 μm). **(c-d)** The number **(c)** and percentage **(d)** of detached *MYC* and *FGFR2* ecDNA per SNU16 metaphase cell after Cyclin C or *CDK8* RNAi knockdown (Kruskal-Wallis test with post-hoc Dunn test). Nocodazole treatment: 12-16 hr. Data were pooled from three replicates for statistics. dAUC was shown as mean ± SEM. **(e)** Representative *MYC* ecDNA FISH on MSTO211H metaphase cells after Cyclin C knockdown (scale bar: 5 μm). **(f-g)** The number **(f)** and percentage **(g)** of detached *MYC* ecDNAs per MSTO211H metaphase cell after Cyclin C RNAi knockdown (two-sided Wilcoxon test). Nocodazole treatment: 8-12 hr. Data were pooled from three replicates for statistics. dAUC was shown as mean ± SEM.

**Extended Data Fig. 7.**
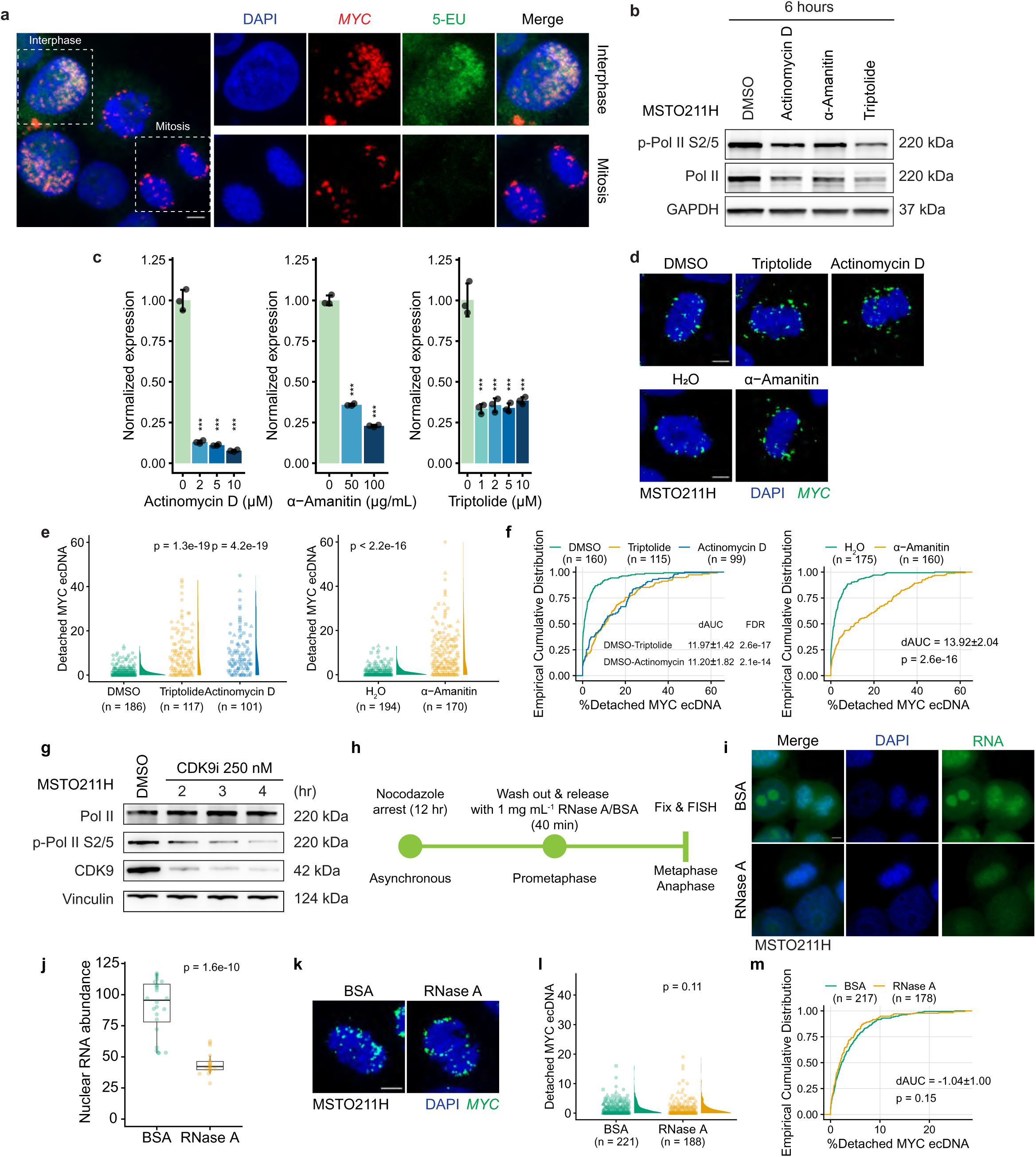
Mitotic transcription and RNA dependency for ecDNA segregation. **(a)** Representative 5-EU labelling images for newly synthesised RNA in interphase and anaphase MSTO211H cells (scale bar: 5 μm). Cells were arrested with nocodazole for 7 hr and were subsequently labelled with 1 mM of 5-EU for 1 hr. **(b)** Western blot for total Pol II and p-Pol II Ser2/5 in MSTO211H cells after transcription inhibitor treatments. **(c)** qPCR results for *MYC* mRNA level in MSTO211H cells after transcription inhibitor treatments. The long half-life *RPL13A* transcript was used for normalisation (one-way ANOVA with post-hoc Dunnett test). **(d)** Representative *MYC* ecDNA FISH on MSTO211H metaphase cells after 2 μM actinomycin D, 2 μM triptolide, or 100 μg mL^-1^ α-amanitin treatment (scale bar: 5 μm). **(e-f)** The number **(e)** and percentage **(f)** of detached *MYC* ecDNAs per MSTO211H metaphase cell after actinomycin D, triptolide (Kruskal-Wallis test with post-hoc Dunn test), or α-amanitin treatment (two-sided Wilcoxon test). Nocodazole treatment: 12 hr. Transcription inhibitors were applied for the last 6 hr during arrest. Data were pooled from three replicates for statistics. dAUC was shown as mean ± SEM. **(g)** Western blot showing a time-series of Pol II, p-Pol II Ser2/5, and CDK9 after 250 nM CDK9i treatment in MSTO211H cells. **(h)** Workflow showing the imaging approach to study ecDNA attachment with mitotic chromosomes during metaphase. A 12-hr nocodazole treatment was first applied to enrich prometaphase cells. After washing out nocodazole, RNase A was added to the cell culture along with releasing cells into metaphase and anaphase for fixation and imaging. **(i)** Representative images for RNA dye signal, which represents cellular RNA level, in MSTO211H cells after 1 mg mL^-1^ RNase A treatment (scale bar: 5 μm). **(j)** Decreased nuclear RNA abundance after 1 mg mL^-1^ RNase A treatment (two-sided Wilcoxon test). **(k)** Representative *MYC* ecDNA FISH in MSTO211H metaphase cells after 1 mg mL^-1^ RNase A treatment (scale bar: 5 μm). **(l-m)** The number **(l)** and percentage **(m)** of detached *MYC* ecDNA per MSTO211H metaphase cell after RNase A treatment (two-sided Wilcoxon test). Nocodazole treatment: 8 hr. Data were pooled from three replicates for statistics. dAUC was shown as mean ± SEM.

**Extended Data Fig. 8.**
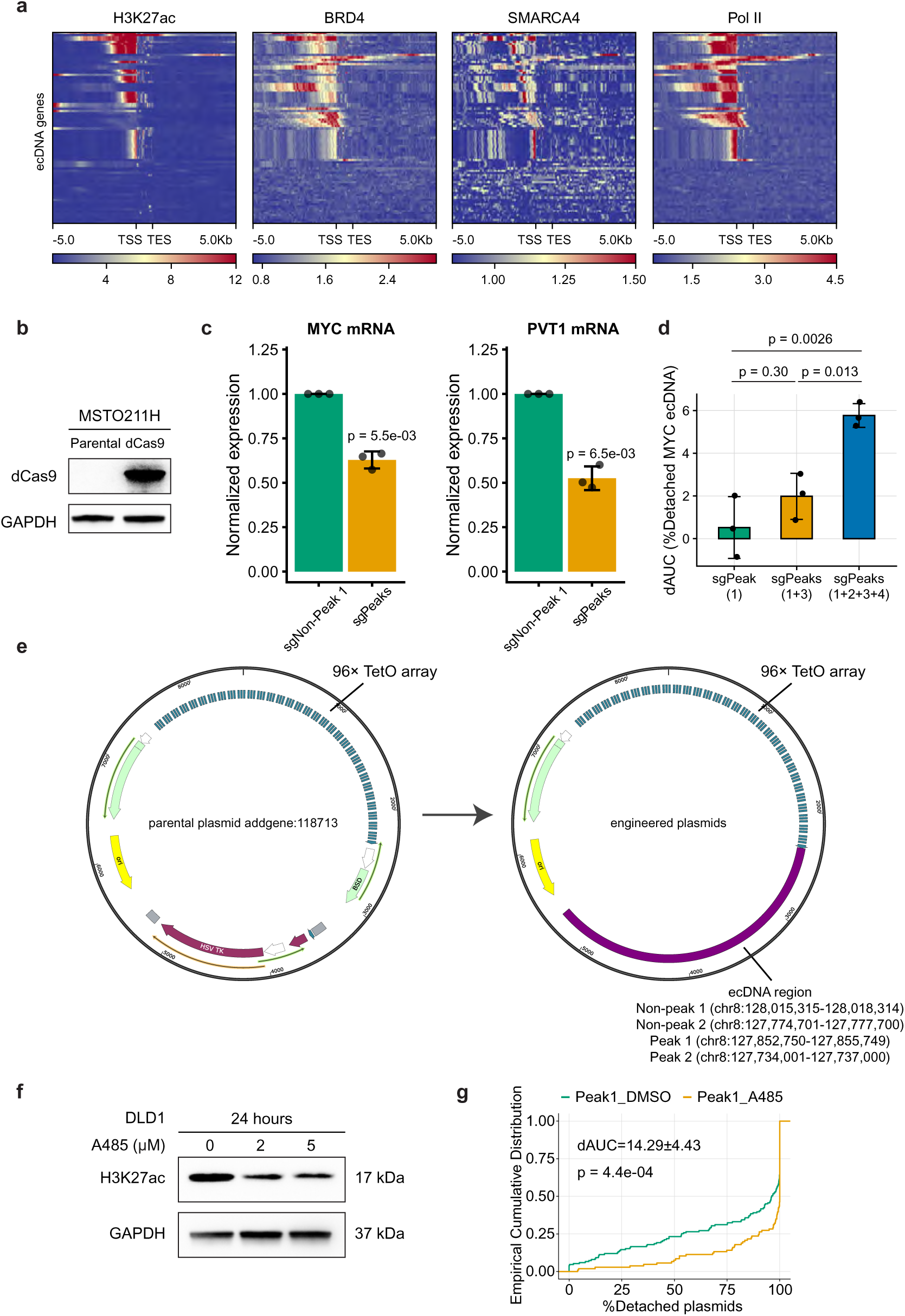
Transcriptionally active ecDNA regions attach plasmids to mitotic chromosomes. **(a)** Binding patterns for H3K27ac, BRD4, SMARCA4, and Pol II in mitotic MSTO211H cells. **(b)** Western blot for dCas9 proteins in parental MSTO211H or MSTO211H-dCas9 cells. **(c)** qPCR results for *MYC* and *PVT1* mRNA levels in MSTO211H cells after CRISPRi. The *TBP* transcript was used for normalisation (two-sided Student’s t-test). **(d)** dAUC comparison when targeting one, two, or four peak regions using CRISPRi (one-way ANOVA with post-hoc Dunnett test). **(e)** The imaging plasmid containing 96× TetO repeats (Addgene 118713, left panel) was engineered to contain transcriptionally inactive (non-peak) or active (peak) ecDNA regions (right panel). **(f)** Western blot for H3K27ac in DLD1 cells after A485 treatment. **(g)** Decreased percentage of plasmids carrying the peak region attached to mitotic chromosomes when the cells were treated with 5 μM A485 (two-sided Wilcoxon test). Data were pooled from three replicates for statistics.

**Extended Data Fig. 9.**
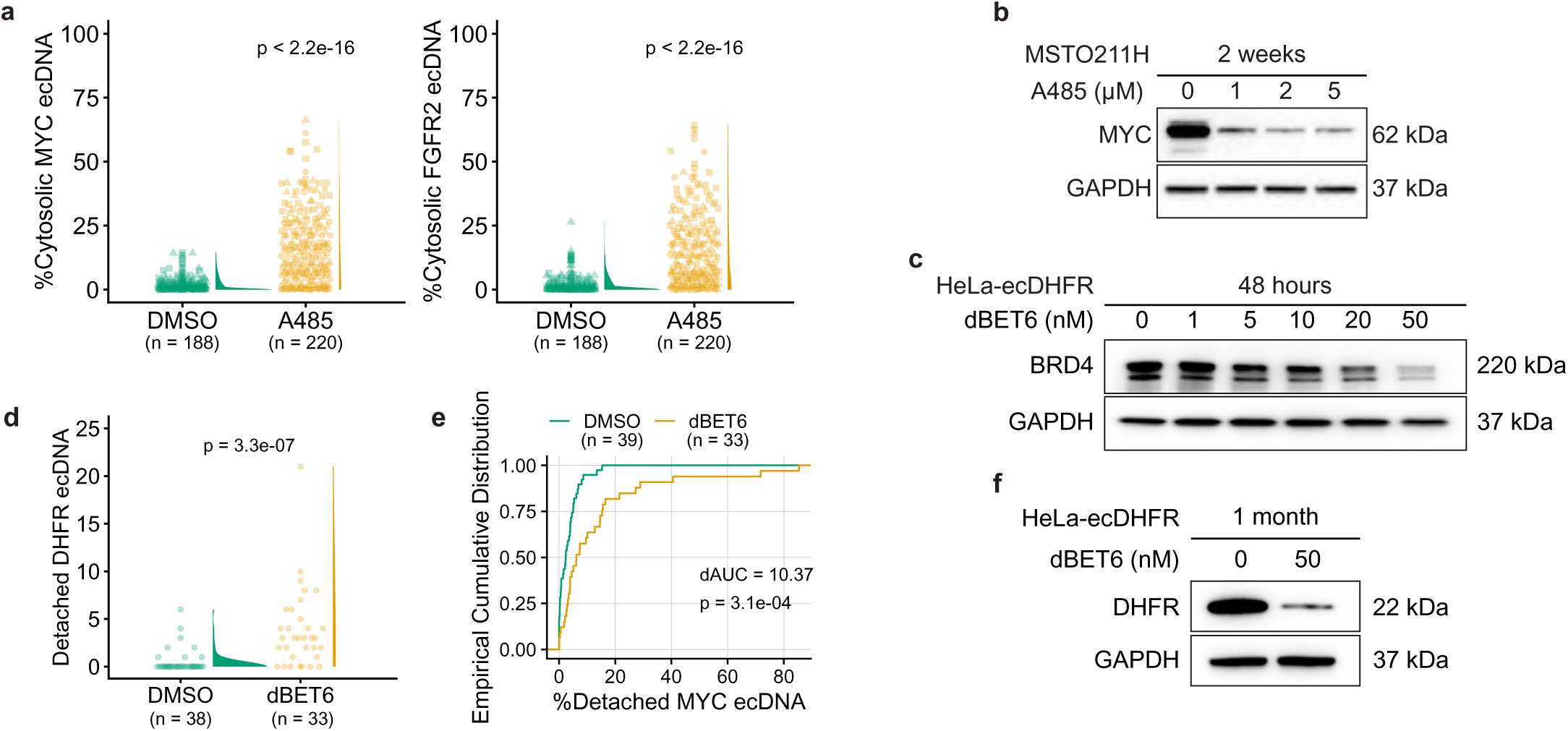
Mis-segregation of ecDNAs reduces oncogenicity. **(a)** Quantification of cytosolic ecDNA content in SNU16 interphase cells (two-sided Wilcoxon test). Cells were treated with 10 μM A485 with a 12-hr nocodazole arrest and were subsequently released for 5 hr to the next cell cycle. Data were pooled from three replicates for statistics. **(b)** Western blot for MYC in MSTO211H cells after 2 weeks of A485 treatment. **(c)** Western blot for BRD4 in HeLa-ecDHFR cells after dBET6 treatment. **(d-e)** The number **(d)** and percentage **(e)** of detached *DHFR* ecDNAs per HeLa-ecDHFR metaphase cell after 48-hr dBET6 treatment (two-sided Wilcoxon test). Nocodazole treatment: 24 hr. Three biological replicates were performed, and one representative is shown. **(f)** Western blot for DHFR in HeLa-ecDHFR cells after 1 month of dBET6 treatment.

